# RNA-mediated MYC multimerization suppresses innate immune signaling

**DOI:** 10.1101/2024.12.06.627168

**Authors:** Leonie Uhl, Amel Aziba, Sinah Löbbert, Timothy Russell, Toshitha Kannan, Omkar R. Valanju, Christina Schülein-Völk, Tim de Martines, Francisco Montesinos, Michael Bolz, Giacomo Cossa, Theresa Endres, Daniel Solvie, Bastian Krenz, Hans M. Maric, Dimitrios Papadopoulos, Seychelle M. Vos, Martin Eilers

**Author notes:** These authors contributed equally: Leonie Uhl and Amel Aziba. Corresponding author: Martin Eilers.

## Abstract

In response to perturbed transcription elongation, the MYC oncoprotein multimerizes and undergoes a phase transition; the underlying mechanisms and their function are unknown. Here, we show that MYC re-localizes from its canonical location on DNA to RNA in response to the accumulation of intronic RNA. MYC binds RNA directly, which enhances its multimerization. MYC multimers concentrate the nuclear exosome, a 3’-5’ RNA exonuclease, and its targeting complexes around double-stranded RNA and R-loops, and promote exosome recruitment to R-loops. RNA binding of MYC suppresses activation of the innate immune kinase TBK1. Upon MYC depletion, intron-derived dsRNAs, including RNA derived from repetitive elements and small nucleolar RNAs, accumulate on TLR3, a pattern recognition receptor that activates TBK1. In MYC-depleted cells, TLR3-bound snoRNAs carry aberrant 3’-ends, indicating defective exosomal processing. Our data show that the phase transition of MYC is a RNA-driven stress response that suppresses the accumulation of immunogenic RNAs.

## Introduction

Deregulated expression of MYC is an oncogenic driver in many tumor entities^1^. The MYC oncoprotein forms a heterodimeric complex with a partner protein MAX, which associates with most active promoters and many enhancers on chromatin^2, 3^. The complex exerts both positive and, in conjunction with additional partner proteins, negative effects on the expression of large sets of genes. Although these effects are weak for individual genes, they result in a characteristic gene expression profile of MYC-driven tumors^2, 3^.

Enhanced expression of MYC exerts multiple cell-autonomous effects: for example, MYC stimulates cell growth and reprograms cellular metabolism, and these effects correlate well with the altered expression of the respective MYC target genes^3^. In addition, MYC proteins have critical non-cell-autonomous functions in oncogenesis, since they establish an immunologically “cold” microenvironment and enable tumors to escape from immune surveillance^4–7^. For example, depletion of endogenous MYC in the KPC model of pancreatic ductual adenocarcinoma (PDAC) causes the elimination of tumors by the immune system^8, 9^. Mechanistically, MYC proteins suppress the expression of MHC class I antigens which present antigens, reducing recognition of tumor cells by T lymphocytes^9, 10^. In addition, MYC suppresses the activity of the TANK-binding kinase 1 (TBK1), which stimulates nuclear factor kappa B (NF-kB) and interferon-dependent genes that attract and activate immune cells. Deletion of TBK1 alleviates the requirement for MYC in PDAC growth^8, 11^. Activation of TBK1 in MYC-depleted cells depends on the TLR3 pattern recognition receptor^8^, which recognizes double-stranded (ds)RNA and R-loops, arguing that MYC suppresses the accumulation of dsRNAs or R-loops on TLR3 and that this is critical for its oncogenic function^12, 13^. Consistently, MYC represses the transcription of genes that are required for TLR3 function^5, 8^ but whether MYC directly affects dsRNA or R-loop metabolism is unknown.

Perturbing transcription elongation by inhibition of the spliceosome or the FACT histone chaperone causes MYC proteins to multimerize and undergo a phase transition^14^, similar to that shown for the neuronal paralog, MYCN^15^. This transition has only small effects on gene expression, but protects replication forks from RNA polymerase to prevent transcription-replication conflicts^14^. While the molecular mechanisms that control MYC multimerization are currently unknown, recent work has revealed that MYC and MYCN can bind to RNA in addition to DNA^16, 17^. Here we show that MYC multimerization reflects the transition from a DNA-to an RNA-bound form of MYC and that MYC multimers suppress TBK1 activation by endogenous dsRNA. Our findings directly link RNA binding and MYC multimerization to its ability to suppress innate immune signaling.

## Results

### MYC broadly binds to nascent RNA in cells

To investigate whether MYC is bound to RNA in cells, we performed cross-linking and immunoprecipitation (CLIP) experiments in K562 chronic myeloid leukemia cells, which, like many tumor cells, express elevated levels of MYC^18^. Cells were exposed to UV-C light, which creates crosslinks between proteins and nucleic acids^19^. After lysis, RNA was fragmented with RNAse I, proteins were immunoprecipitated, and pCp-biotin was ligated to RNA before protein-RNA complexes were separated by SDS-PAGE (Figure 1a). Immunoprecipitations (IPs) of MYC from UV-crosslinked (XL) K562 cells stained positive for RNA, with the signal being sensitive to RNase I. Control IgG IPs and MYC-IPs from cells that were not UV-cross-linked (NXL) contained no RNA, confirming the specificity of the signal. Immunoprecipitated RNA was then prepared for single-end enhanced CLIP (seCLIP) sequencing^19^. Analysis of replicate seCLIPs using the Skipper algorithm^20^ identified 11,077 reproducible enriched windows in MYC-IPs versus 72 reproducible enriched windows in IgG controls (Figure 1b). Heatmaps as well as inspection of individual gene loci confirmed the specificity of the MYC signal over IgG controls (Figure 1c,d). Around 70% of MYC bound sites were localized in intronic RNA, and ∼10% of MYC binding sites were located adjacent to splice sites indicating binding of MYC to nascent or unprocessed transcripts, consistent with findings for MYCN^16^ (Figure 1e). Comparison with ENCODE data showed that the overall distribution of MYC binding sites on RNA was similar to that of other intron-binding proteins such as FUS, MATR3 and PTBP1^21–23^ (Figure 1f and Extended Data Figure 1a). The Skipper algorithm identified multiple MYC-bound windows in many introns, suggesting that introns are potentially bound by multiple MYC molecules (Figure 1g). GO term analysis did not reveal any enrichment of a functional class of transcripts among MYC-bound genes and comparison with input data suggested that MYC can bind to most unprocessed transcripts in a cell. *De novo* motif enrichment analysis of MYC-bound RNA windows revealed a moderate enrichment of sequence motifs bound by MATR3 and PTPB1^24^. Importantly, this did not reveal any preference for E-box-consensus or -related sequences, to which MYC/MAX complexes bind on DNA, arguing that the RNA association of MYC is mechanistically distinct from its ability to bind DNA (Extended Data Figure 1b). To investigate how universally MYC binds to nascent RNA, we also performed seCLIP experiments in U2OS expressing doxycycline-inducible ectopic MYC (U2OS^MYC-Tet-On^ cells). U2OS cells express 1.8×10^5^ endogenous MYC molecules per cell^14^, which is comparable to proliferating non-transformed cells (e.g. IMEC, HMLE, MCF10A) and lower than those found in many tumor cell lines^25^. In EtOH-treated (control) U2OS^MYC-Tet-On^ cells the overall binding of endogenous MYC to RNA was very similar to that observed in K562 cells: MYC bound to thousands of RNAs (Extended Data Figure 1c) with MYC binding being predominantly in introns (Extended Data Figure 1d).

**Figure 1:**
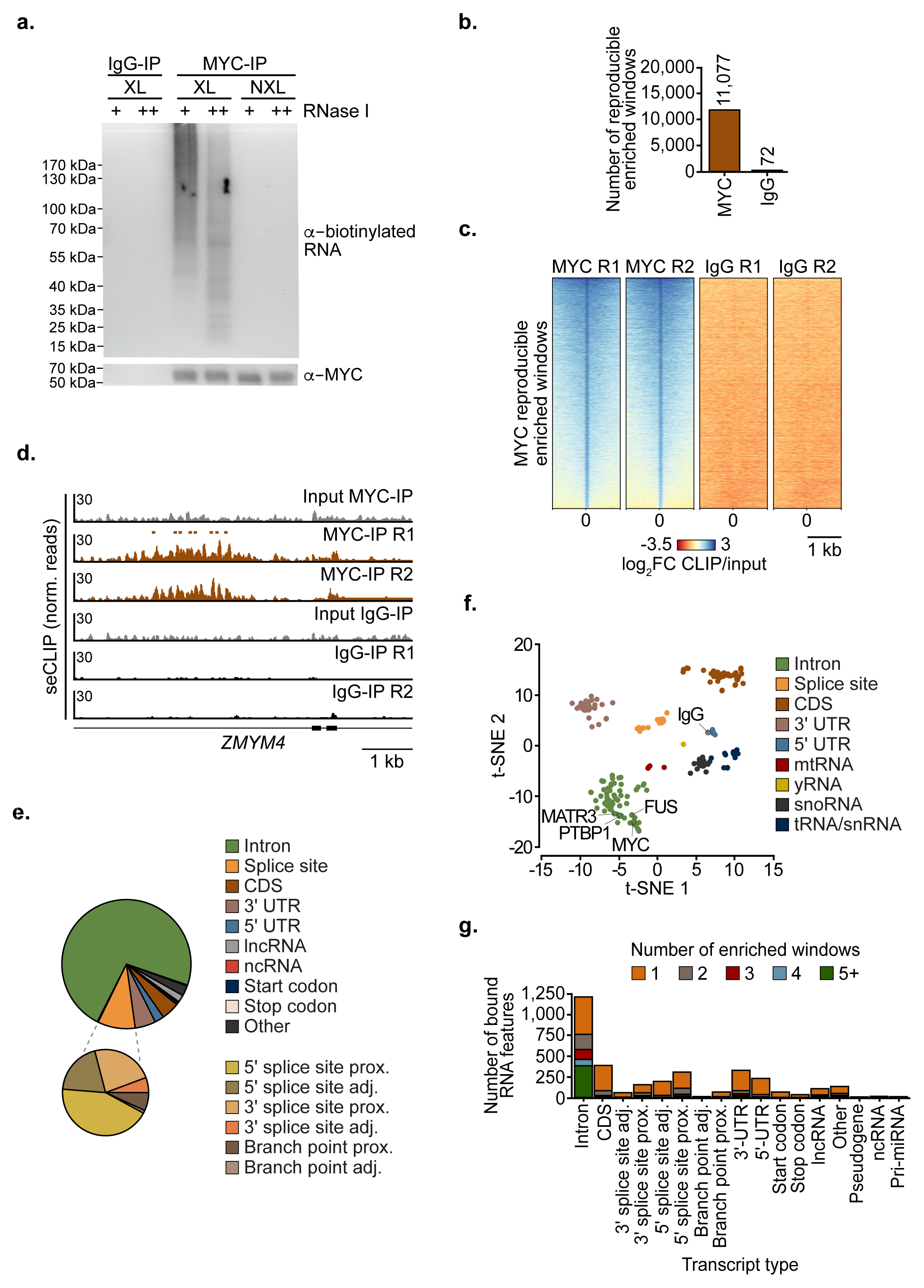
MYC binds to nascent RNA. **a.** Top: Visualization of biotinylated RNA recovered in MYC immunoprecipitations (IPs) from lysates of UV cross-linked (XL) K562 cells treated with standard (40 U; +) or high (333 U; ++) levels of RNase I. Beads coupled with non-specific IgG and non-cross-linked (NXL) K562 cells were used as controls. Bottom: Corresponding α-MYC immunoblot (*n*=2). **b.** Number of reproducible enriched windows in α-MYC and control IgG seCLIPs in K562 cells identified using Skipper (*n*=2). **c.** Heatmap showing log_2_FC CLIP/input at reproducible enriched windows called in two MYC seCLIP replicates from K562 cells. Heatmap sorted by descending log_2_FC CLIP/input means of the MYC R1 seCLIP (*n*=2). **d.** Browser picture of MYC and IgG control seCLIPs at the *ZMYM4* gene. Bars shown above replicate 1 (R1) of the MYC-IP indicate the location of reproducible enriched windows (*n*=2). **e.** Pie chart showing number of reproducible enriched windows stratified for different RNA feature types (*n*=2). f. Skipper t-SNE plot of MYC against ENCODE eCLIP RBP data. Colors indicate the RNA feature preference of the respective classes of RBPs. **g.** Number of RNA features containing reproducible enriched windows in MYC seCLIPs. Colors indicate the number of windows per RNA feature.

### MYC proteins exist in a dynamic equilibrium of DNA- and RNA-bound pools

In response to inhibition of the FACT histone chaperone or the SF3B1 subunit of the spliceosome, MYC proteins undergo a phase transition and multimerize^14^, prompting us to investigate whether inhibition of FACT or the spliceosome affects the distribution of MYC between RNA- and DNA-bound pools. Since exposure to the FACT inhibitor CBL0137 reduced endogenous *MYC* mRNA levels, we performed these experiments in U2OS^MYC-Tet-On^ cells that had been treated with doxycycline for 24 hours. Under these conditions, U2OS^MYC-Tet-On^ cells express approximately 3×10^6^ molecules of MYC per cell, about 3-fold higher than MYC levels in colon carcinoma^25^ and glioblastoma cells^26^. CLIP experiments revealed an increase in MYC RNA binding upon doxycycline-induction that paralleled the increase in MYC protein levels (Extended Data Figure 2a,b). seCLIP sequencing revealed that the overall binding pattern of MYC to RNA was very similar between low and elevated MYC levels (Extended Data Figure 2c). Skipper analyses revealed a twofold increase in the overall number of MYC binding sites on RNA and showed that, while binding to introns was largely conserved between low and elevated MYC levels, MYC showed enhanced binding to exons at elevated levels (Extended Data Figure 2d,e). To determine how FACT blockade affects the distribution of MYC between DNA- and RNA-bound pools, we performed parallel seCLIP sequencing of doxycycline-treated U2OS^MYC-Tet-On^ cells that had additionally been treated for 2 h with CBL0137. Exposure to CBL0137 strongly increased both the number of MYC RNA binding sites and RNA binding at many individual RNAs (Figure 2a,b). Overall, 25% of RNA binding sites of MYC that were identified in unperturbed cells were also enriched in CBL0137-treated cells, with several thousand additional sites appearing in CBL0137-treated cells (Figure 2c). The majority of these new binding sites were intronic, suggesting that the induction of transcriptional stress increases MYC occupancy at unprocessed transcripts (Figure 2d). Previous chromatin-immunoprecipitation (ChIP) assays had shown a decrease in MYC binding to several individual E-box containing promoters upon FACT inhibition^14^. ChIP-Rx sequencing from doxycycline-treated U2OS^MYC-Tet-On^ cells showed that FACT inhibition caused a virtually complete loss of MYC from active promoters (Figure 2e). Similarly, inhibition of SF3B1 by pladienolide B (PlaB) caused a global redistribution of MYC from active promoters to intronic RNA with an increased proximity to splice sites (Extended Data Figure 2f-h). Since inhibition of FACT or the spliceosome cause an increase in unprocessed RNAs^27, 28^, we hypothesized that the increased RNA binding of MYC reflects the accumulation of intronic RNAs during perturbed transcription elongation. To test this, we used siRNAs to deplete EXOSC5, a core subunit of the nuclear exosome, which degrades intronic RNAs^29^. Inspection of both individual genes (Figure 2f) and average density plots (Figure 2g) of MYC seCLIP experiments revealed that depletion of EXOSC5 caused an increase in MYC RNA binding, which was reflected in the increased total levels of mainly intronic RNAs. Parallel experiments performed in cells treated with siRNA targeting MYC confirmed the specificity of the signal (Figure 2f,g). We concluded that MYC exists in a dynamic equilibrium between DNA- and RNA-bound pools and that accumulation of intronic RNAs shifts MYC from its canonical DNA-bound to an RNA-bound state.

**Figure 2:**
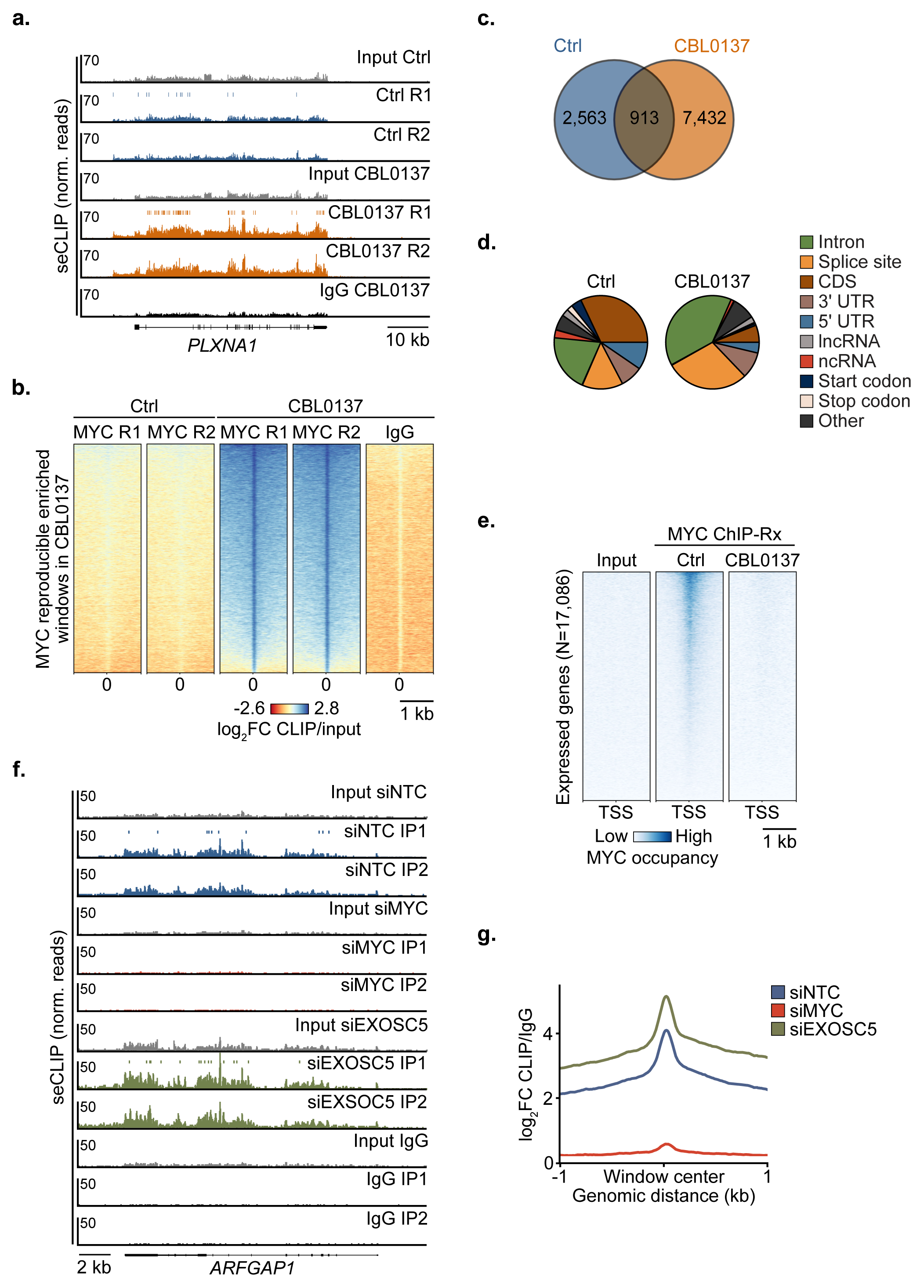
RNA and DNA binding of MYC are in a dynamic equilibrium. **a.** Browser picture of MYC and IgG control seCLIPs at the *PLXNA1* gene. Cells were treated with CBL0137 (5 µM, 2 h) or DMSO as control. Bars above Ctrl R1 and CBL0137 R1 MYC seCLIPs indicate the position of reproducible enriched windows (*n*=2). **b.** Heatmap showing the log_2_FC CLIP/input of MYC seCLIPs at reproducible enriched windows called upon treatment with CBL0137 (5 µM, 2 h) in U2OS^MYC-Tet-On^ cells treated with doxycycline (1 µg/mL, 24 h). Heatmap sorted by descending log_2_FC CLIP/input means the MYC R1 seCLIP upon CBL0137 treatment (*n*=2). **c.** Venn diagram showing the overlap between reproducible enriched windows called in control and CBL0137-treated cells. Treatment conditions as described in a (*n*=2). **d.** Pie charts showing the distribution of RNA classes of reproducibly enriched windows (*n*=2). **e.** Heatmap of MYC chromatin occupancy at the transcription start site (TSS) of all expressed genes analyzed by ChIP-Rx sequencing in U2OS^MYC-Tet-On^ cells treated with doxycycline (1 µg/mL, 24 h) and CBL0137 (5 µM, 2 h) or DMSO as control. Heatmap sorted by descending mean reads in the Ctrl condition (*n*=2). **f.** Exemplary browser pictures of MYC seCLIPs at the *ARFGAP1* gene in U2OS^MYC-Tet-On^ cells treated with doxycycline (1 µg/mL, 24 h) and transfected with siEXOSC5 or a non-targeting control (siNTC). Cells transfected with siMYC and IPs using non-specific IgG antibodies were used as controls (*n*=2). **g.** Average density plot showing the log_2_FC CLIP/IgG of MYC seCLIPs at reproducible enriched windows called in MYC seCLIPs transfected with either siNTC or siEXOSC5 under conditions described in f.

### MYC multimers co-localize with RNA degrading factors around R-loops and dsRNA

Proximity labeling experiments had indicated a dramatic change in the MYC interactome upon multimerization and shown that MYC multimers are localized close to many RNA processing enzymes^14^. To explore whether MYC associates with RNA metabolizing or degrading factors in unstressed cells, we performed IPs from control U2OS^MYC-Tet-On^ cells (Figure 3a). Consistent with previous data, IPs of endogenous MYC co-immunoprecipitated MAX as well as total and S2-phosphorylated (elongating) RNA Polymerase II (RNAPIIpS2) (Figure 3a). Anti-MYC, but not control IPs, also contained EXOSC10, a catalytic subunit of the exosome, ZCCHC8, the scaffolding subunit of the nuclear exosome targeting complex (NEXT), YTHDC1, RBM15 and METTL3, all of which are involved in m6A metabolism, as well as the RNA binding proteins FUS, HNRNPC and STRBP (Figure 3a). In these experiments, we added an RNase inhibitor to the lysate. To rule out that these interactions were mediated indirectly via binding to RNA, we showed that treatment of cell lysates with benzonase, which degrades both DNA and RNA, had no effect on the interaction of MYC with multiple RNA degrading factors (Extended Data Figure 3a). We confirmed the interactions of MYC with several RNA processing factors by reverse co-IPs (Extended Data Figure 3b). To understand whether the interaction of MYC with RNA metabolizing factors is constitutive or stress-regulated, we repeated the immune precipitations after treatment of cells with MG-132 or CBL0137, both of which induce MYC multimerization. This confirmed that MYC associates with multiple RNA metabolizing and degrading factors in unstressed cells and revealed that both treatments caused changes in the amount of co-immunoprecipitating proteins that paralleled overall changes MYC levels, arguing that MYC binds to these partner proteins constitutively (Extended Data Figure 3c). We then performed quantitative immunofluorescence experiments from doxycycline-treated U2OS^MYC-Tet-On^ cells that had been exposed to CBL0137 and found that multiple factors involved in RNA metabolism and degradation were concentrated in MYC multimers (Figure 3b). This was specific, since parallel experiments with several transcription termination factors did not reveal any co-localization with MYC multimers (Extended Data Figure 3d).

**Figure 3:**
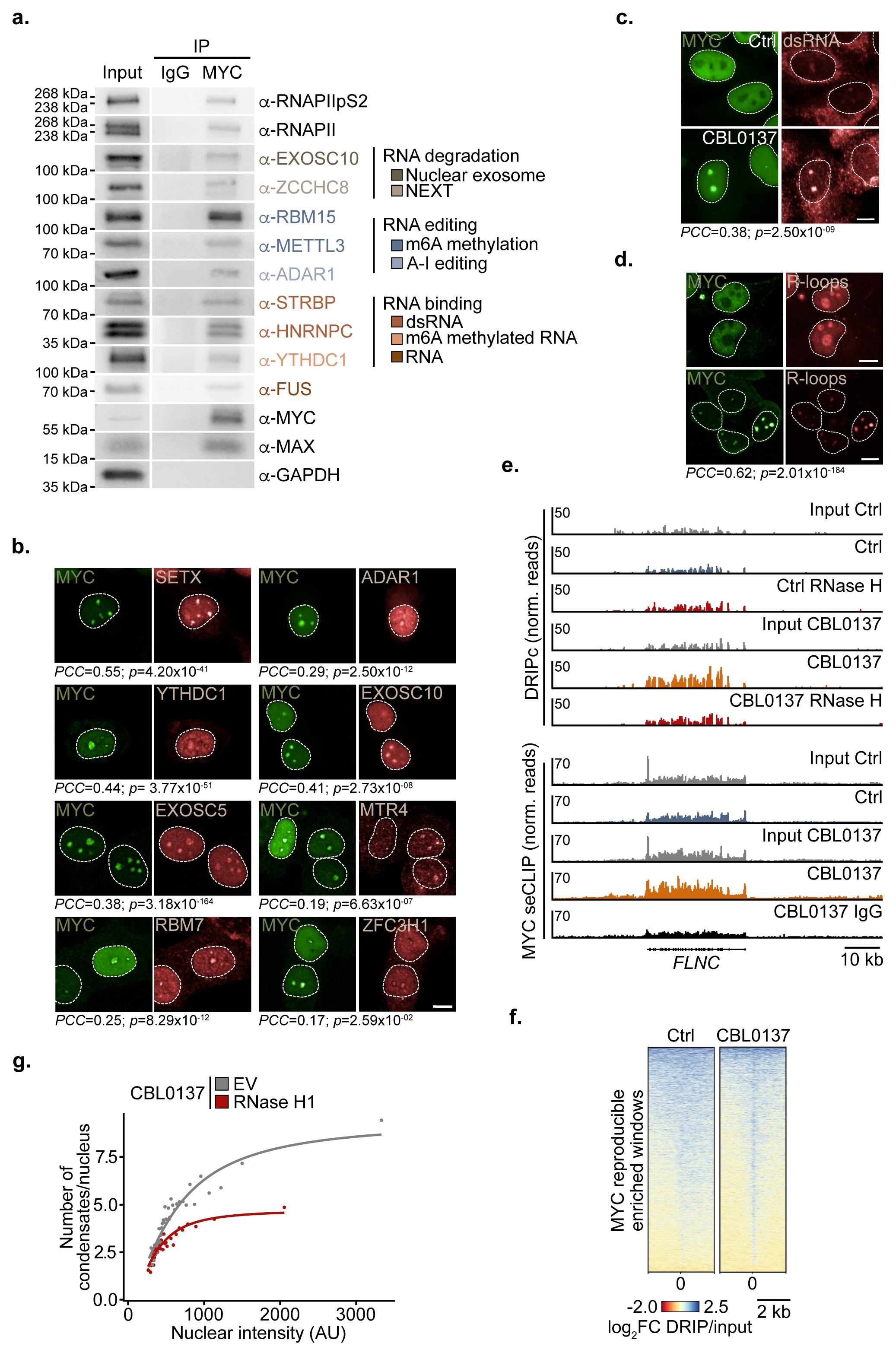
MYC multimers are hubs of RNA degrading factors. **a.** Immunoblot of MYC-IP from U2OS^MYC-Tet-On^ EtOH-treated control cells. Samples were incubated with RNase inhibitor (50 U/mL) before precipitation. Colours of the RNA processing factors indicate functional categories of RNA binding complexes as indicated. Input is 1% for all IPs unless otherwise indicated. GAPDH was used as loading control (*n*=3). **b.** Representative immunofluorescence of indicated proteins in U2OS^MYC-Tet-On^ cells treated with doxycycline (1 µg/mL, 24 h) and CBL0137 (5 µM, 4 h). Scale bar: 10 µm (*n*=3). Pearson Correlation coefficients (*PCCs*) between the partitioning ratio of MYC and of the protein of interest (POI) are indicated below the respective immunofluorescence pictures. The *p*-value is the significance level of Pearson’s product-moment correlation *t*-test, two-sided (*n*=3). **c.** Immunofluorescence of MYC and double-stranded (ds)RNA (J2 antibody) in U2OS^MYC-Tet-On^ cells treated with doxycycline (1 µg/mL, 24 h) and CBL0137 (5 µM, 4 h) or DMSO as control (*n*=5). The co-localization was quantified in CBL-treated cells as described in b. **d.** Immunofluorescence of MYC and R-loops in U2OS^MYC-Tet-On^ cells that stably express the hybrid binding domain (HBD) of RNase H1 fused to GFP treated with doxycycline (1 µg/mL, 24 h) and CBL0137 (5 µM, 4 h) or DMSO as control. Scale bar: 10 µm (*n*=5). The co-localization was quantified in CBL-treated cells as described in b. **e.** Exemplary browser pictures of spike normalized DRIPc and MYC seCLIPs in U2OS^MYC-Tet-On^ treated with doxycycline (1 µg/mL, 24 h) and CBL0137 (5 µM, 2 h) or DMSO as control at the *FLNC* gene (DRIPc, *n*=1; MYC seCLIP *n*=2, shown is one representative replicate). **f.** Heatmap of R-loops as determined by spike-normalized DRIPc sequencing at MYC RNA-binding sites in control and CBL0137-treated U2OS^MYC-Tet-On^ cells (*n*=1). **g.** Number of condensates per nucleus in EV and NLS-RNase H1-FLAG expressing U2OS^MYC-Tet-On^ cells treated with doxycycline (1 µg/mL, 24 h) and CBL0137 (5 µM, 4 h) relative to the binned nuclear MYC intensity. 3921 and 3101 cells were analyzed for the EV and RNase H1 condition respectively, and divided into 40 bins (*n*=8).

To determine whether specific RNA structures are localized within MYC multimers, we stained cells with J2, a monoclonal antibody that recognizes double-stranded (ds)RNA. In addition, we stably expressed the hybrid-binding domain (HBD) of RNase H1, which recognizes DNA/RNA-hybrids, fused to an antibody tag. To validate the specificity of R-loop staining, we used a mutant allele (“WKK”) that carries three point mutations in the HBD (“WKK”) (Extended Data Figure 4a). Staining of doxycycline-treated U2OS^MYC-Tet-On^ cells showed that both dsRNA and R-loops were concentrated in MYC multimers in the presence of CBL0137 (Figure 3c,d). To extend these observations, we performed DNA:RNA immunoprecipitation followed by cDNA conversion (DRIPc) sequencing which maps the genome-wide distribution of R-loops (Extended Data Figure 4b, c). Strinkingly, there was no significant enrichment of R-loops at MYC-bound RNAs in control cells, while R-loops identified in CBL0137-treated cells overlapped with MYC binding sites on RNA (Figure 3e, f). To determine whether the presence of R-loops is required for MYC multimerization, we stably expressed RNase H1 fused to a nuclear-localization signal in U2OS^MYC-Tet-On^ cells and confirmed that the enzyme localizes to the nucleus (Extended Data Figure 4d). Parallel measurements of MYC levels and number of multimeric foci showed that expression of RNase H1 attenuated MYC multimerization in response to CBL0137 treatment; to exclude that this decrease was an indirect consequence of changes in MYC levels, we quantified the MYC immunofluorescence signal and the number of foci independently in more than 500 cells and plotted the number of foci in bins of cells with closely similar MYC levels (Figure 3g and Extended Data Figure 4e). We concluded that MYC multimers form in hubs that concentrate RNA degrading, but not transcription termination factors, in the vicinity of R-loops and double-stranded RNA and that R-loops promote multimerization of MYC.

### Failure to degrade nascent RNA drives MYC multimerization

To pinpoint the factors that drive MYC multimerization, we performed an siRNA screen that targeted 84 proteins that we identified in previous analyses of the interactome of MYC multimers^14^ or of RNA-bound MYCN^16^ (Extended Data Figure 4f). We transfected U2OS^MYC–Tet-On^ cells that had been treated with doxycycline to minimize effects on MYC expression with specific siRNAs and measured their effect on MYC multimer formation in unstressed cells by high-content microscopy. As controls we confirmed that siRNAs targeting *SUPTH16* and *SSRP1*, two subunits of the FACT complex targeted by CBL0137, and *SF3B1*, a subunit of the spliceosome, promoted MYC multimerization (Extended Data Figure 4g)^14^. We found that several siRNAs induced MYC multimerization in unstressed cells: these targeted predominantly RNA processing and degrading factors and proteins involved in m6A metabolism (Figure 4a,b). To further define the mechanisms that control MYC multimerization, we performed a second siRNA screen targeting specifically subunits of the nuclear exosome and its targeting complexes, TRAMP, NEXT, and PAXT, as well as additional factors, such as ARS2, that determines early transcript fate^30^. This showed that depletion of subunits of the exosome as well as the NEXT complex, that targets the exosome to degrade non-polyadenylated RNAs, triggers MYC multimerization, whereas depletion of other factors had weak or no effects (Figure 4c,d). Upon FACT inhibition, MYC multimers are often found close to stalled replication forks, which are characterized by the presence of FANCD2, phosphorylated ATR and accumulation of the single-strand binding protein RPA^14^. We confirmed that the same occurs upon depletion of several exosome subunits, the NEXT subunit ZCCHC8 and other RNA degrading factors (Figure 4e and Extended Data Figure 4h). We also noted that MYC binding sites on RNA overlapped to a significant degree with those of EXOSC10 (Figure 4f). We concluded that a failure to degrade nascent or aberrant RNAs drives MYC multimerization.

**Figure 4:**
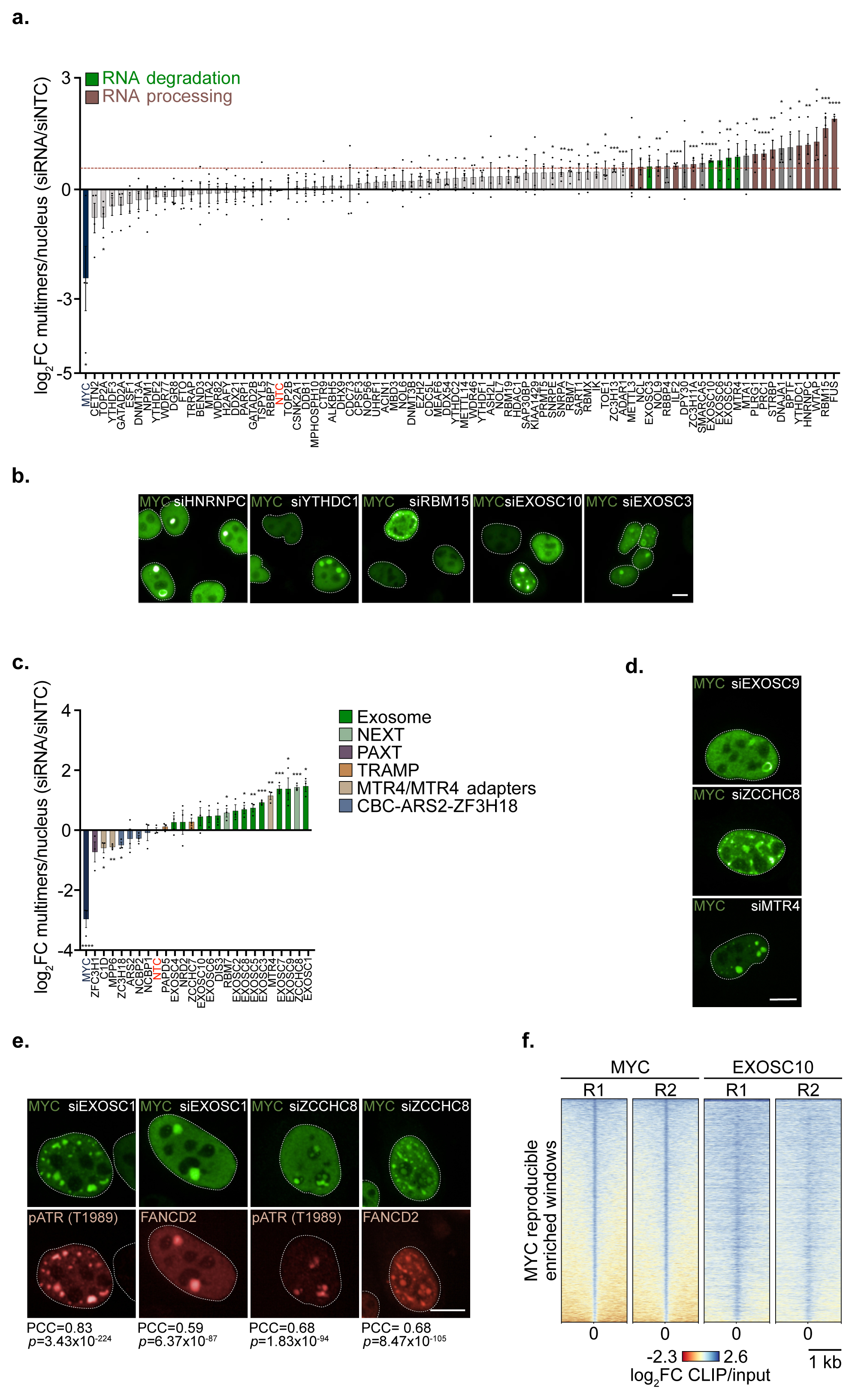
Perturbation of RNA processing drives MYC multimerization. **a.** Targeted siRNA screen quantifying the number of MYC multimer foci per cell after transfection of the indicated siRNAs (48 h) into U2OS^MYC-Tet-On^ cells treated with doxycycline (1 µg/mL, 24 h). Dashed red line indicates a log_2_FC of 0.58. Above this threshold, siRNAs were categorized into RNA degradation or processing factors and coloured as indicated. The non-targeting control (NTC) is colored in red, siMYC is marked in dark blue. Means of biological replicates (*n=*4) ± s.e.m. are shown. Asterisks indicate *p*-values <0.05 (*), <0.01 (**), <0.001 (***), <0.0001 (****), unpaired, two-sided *t*-test. **b.** Representative immunofluorescence images of MYC from targeted siRNA screen shown in Scale bar: 10 µm (*n*=4). **c.** Targeted siRNA screen quantifying the number of MYC multimer foci per cell after transfection of the indicated siRNAs (48 h) into U2OS^MYC-Tet-On^ cells treated with doxycycline (1 µg/mL, 24 h). The non-targeting control (NTC) is colored in red, siMYC is marked in dark blue. Means of biological replicates (*n=*3) ± s.e.m. are shown. Asterisks indicate *p*-values <0.05 (*), <0.01 (**), <0.001 (***), <0.0001 (****), unpaired, two-sided *t*-test. **d.** Representative immunofluorescence images of MYC from targeted siRNA screen shown in Scale bar: 10 µm (*n*=3). **e.** Immunofluorescence images of MYC, FANCD2 and pATR (T1989) in U2OS^MYC-Tet-On^ treated with doxycycline (1 µg/mL, 24 h) and indicated siRNAs (48 h). Scale bar: 10 µm (*n=*3). The co-localization was quantified using *PCC*, *t-*test, two-sided. **f.** Heatmap of the log_2_FC CLIP/input of MYC seCLIPs in U2OS^MYC-Tet-On^ cells treated with doxycycline (1 µg/ml, 24 h) overlaid with the log_2_FC CLIP/input of publicly available EXOSC10 eCLIP data (ENCODE) (*n*=2).

### MYC binds directly to RNA and recruits EXOSC10 to R-loops

These findings suggest that MYC multimers represent the RNA-bound form of MYC. To test this hypothesis, we used peptide libraries consisting of overlapping 25mers peptides spanning the entire MYC sequence to identify the residues that mediate RNA binding. This analysis identified four regions (“RBR”: <u>R</u>NA <u>b</u>inding regions), capable of RNA binding, with a domain surrounding MYCBox IV^31^ showing the strongest binding. In addition, sequences surrounding MYCBox I, close to MYCBox II, and the DNA-interacting basic region (BR) were able to bind RNA (Figure 5a). The BR contains an arginine-rich RNA-binding motif recently identified in many transcription factors^17^; we did not investigate the BR further as mutations in the BR would also disrupt DNA binding^32^. In fluorescence polarization assays, recombinant proteins encompassing amino acids 1-163 and amino acids 304-354 showed significant affinity to an RNA oligonucleotide that has been designed based on an RNA sequence robustly bound by MYC in multiple seCLIP experiments (Figure 5d, Extended Data Figure 5a)^16^. Since the RBRs overlap with sequences involved in protein modification and turnover, we first stably expressed several alleles in which basic residues in these domains had been replaced by alanine (Extended Data Figure 5b) in U2OS cells, in which the endogenous MYC gene has been tagged with an auxin-inducible degron (U2OS^MYC–AID^ cells)^14^. All amino-terminal replacement mutants were expressed at significantly lower levels than wildtype MYC (WT MYC), precluding their further analysis in cells (Figure 5c, Extended Data Figure 5c). In contrast, a mutant allele carrying six alanine replacements in the RNA binding domain surrounding MYCBox IV (K316A, K317A, R331A, R334A, R340A, K341A) was expressed at the same level as wildtype MYC (Figure 5b,c). *In vitro* binding assays confirmed that these mutations abolished RNA binding to an oligonucleotide from one of the most strongly MYC bound sites that is located in an intron of the PTDSS2 gene (Figure 5d), and we therefore performed subsequent assays with this allele (“RBRIII^MUT^ MYC”). Notably, RBRIII^MUT^ MYC retains the HCF-1 binding site^31^ in MYCBox IV and three basic amino acids (K323, R324, K326) that are part of the nuclear localization signal and was exclusively localized in the nucleus (Figure 5b,e)^33^. Relative to WT MYC, RBRIII^MUT^ MYC was impaired in its ability to form multimers (Figure 5f, h) and and to co-localize with EXOSC10 upon CBL0317 treatment (Figure 5g,h). Notably, PLA assays showed that the proximity of EXOSC10 to R-loops was significantly decreased in cells expressing RBRIII^MUT^ MYC relative to cells expressing WT MYC (Figure 5i,j), arguing that RNA-bound MYC recruits the nuclear exosome to R-loops.

**Figure 5:**
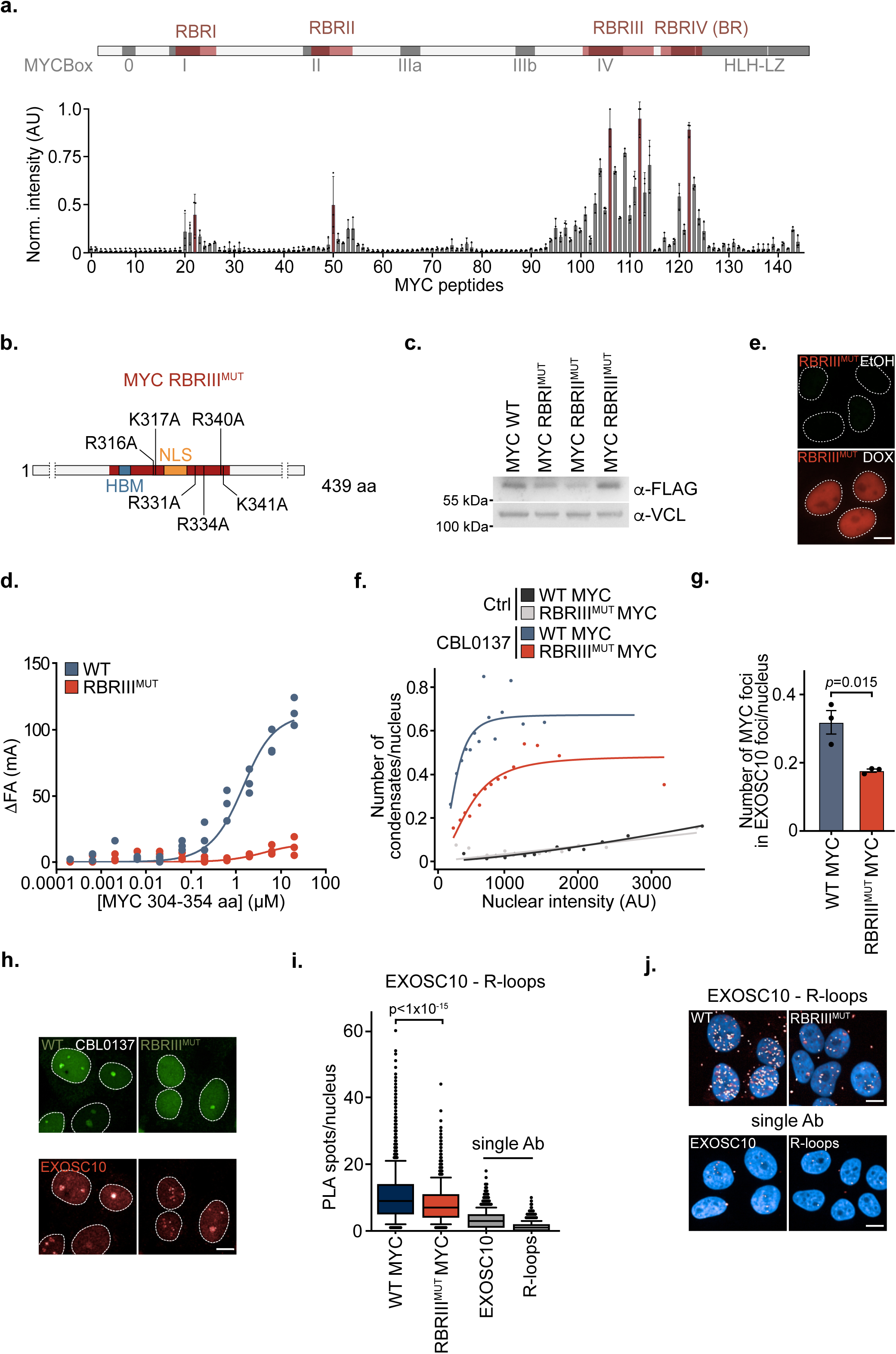
MYC binds directly to RNA. **a.** Peptide microarray of MYC (aa 1-454) displayed as 144 overlapping 25mer peptides with an offset of 3 amino acids. Arrays were incubated with Cy5-labeled RNA. Data show mean ± s.d. of fluorescence intensities normalized to the maximum intensity measured (*n=*3). The MYC sequence is superimposed on the regions corresponding to the respective peptides. RBR: <u>R</u>NA <u>B</u>inding <u>R</u>egion. Conserved sequence elements are shown in grey. **b.** Graphical overview of alanine mutations in MYC in the RNA binding region III indicating that basic amino acids of the nuclear localization sequence (NLS) and HCF-1 binding domain (HBM) remain unaltered. **c.** Immunoblot of U2OS^MYC-AID^ cells stably expressing doxycycline-inducible FLAG-tagged wildtype MYC (WT MYC) or mutants. Cells were treated with doxycycline (1 µg/mL, 24 h) and IAA (500 µM, 24 h). Vinculin (VCL) is shown as loading control (*n*=2). **d.** Fluorescence anisotropy (FA) measurement using half-log dilutions from 20 μM to 0.2 nM of WT or RBR^MUTIII^ MYC constructs (304-354 aa) with an RNA oligonucleotide from one of the strongest MYC bound RNAs (PTDSS2). Three biological replicates are shown for each dilution (*n*=3). **e.** Immunofluorescence of RBRIII^MUT^ MYC expressing U2OS^MYC-AID^ cells treated with doxycycline (1 µg/mL, 24 h) or EtOH as control. Scale bar: 10 µm (*n*=3). **f.** Number of MYC condensates/nucleus in WT MYC and RBRIII^MUT^ MYC expressing U2OS^MYC-AID^ cells treated with doxycycline (1 µg/mL, 24 h), IAA (500 µM, 24 h) and CBL0137 (5 µM, 4 h) relative to the nuclear MYC intensity. Cells were divided into 15 bins based on the nuclear MYC intensity. Number of cells analyzed per condition; DMSO WT MYC: 5,486, CBL0137 WT MYC: 3,326, DMSO RBRIII^MUT^ MYC: 6,431, CBL0137 RBRIII^MUT^ MYC: 4,947 (*n*=4). **g.** Quantification of MYC condensates in EXOSC10 spots/nucleus in WT MYC and RBRIII^MUT^ MYC expressing U2OS^MYC-AID^ cells treated as indicated in f and g. Shown is the mean of biological replicates ± s.e.m., unpaired, two-sided *t*-test (*n*=3). **h.** Representative images of WT MYC and RBRIII^MUT^ MYC expressing U2OS^MYC-AID^ cells treated as described in f. Cells were co-stained for EXOSC10. Scale bar: 10 µm (*n*=3). **i.** Box plot of a proximity ligation assay (PLA) between EXOSC10 and R-loops in U2OS^MYC-Tet-On^ cells that stably express the hybrid binding domain of RNase H1 fused to GFP treated with doxycycline. The box plot depicts the median and the range between the 25th and 75th percentiles. The whiskers are displayed down to the 10th and up to the 90th percentile, and points outside this range are shown as individual dots. Two-sided Mann-Whitney-U test was used for comparison between WT MYC and RBRIII^MUT^ MYC. PLA data were analyzed with the following number of cells: WT: 4,318, RBRIII^MUT^: 2,874 over three independent experiments (*n*=3). **j.** Representative images from PLA between EXOSC10 and R-loops in U2OS^MYC-Tet-On^ cells quantified in i. Scale bar: 10 µm (*n*=3).

### RNA binding of MYC is required to suppress innate immune signaling

To explore the biological function of MYC RNA binding, we expressed WT MYC and RBRIII^MUT^ MYC in murine pancreatic cancer cells that are driven by mutant KRAS^G12D^ and Tp53^R172H^ (KPC cells)^34^ and that express doxycycline-inducible shRNAs targeting endogenous MYC (Figure 6a)^8, 9^. Immunoblots confirmed the depletion of endogenous MYC upon doxycycline addition and also confirmed that WT MYC and RBRIII^MUT^ MYC were expressed at equal levels (Figure 6a). Expression of WT MYC and, albeit to a lesser degree, RBRIII^MUT^ MYC stimulated the proliferation of KPC cells (Figure 6b). RNA sequencing from cells treated for 48 h with doxycycline showed that WT MYC regulated characteristic target gene sets encoding, for example, proteins involved in protein translation, consistent with many previous observations (Figure 6c). In these experiments, RBRIII^MUT^ MYC was indistinguishable in its effects on gene expression from WT MYC (Figure 6c), arguing that RNA binding has no effect on transcriptional regulation by MYC.

**Figure 6:**
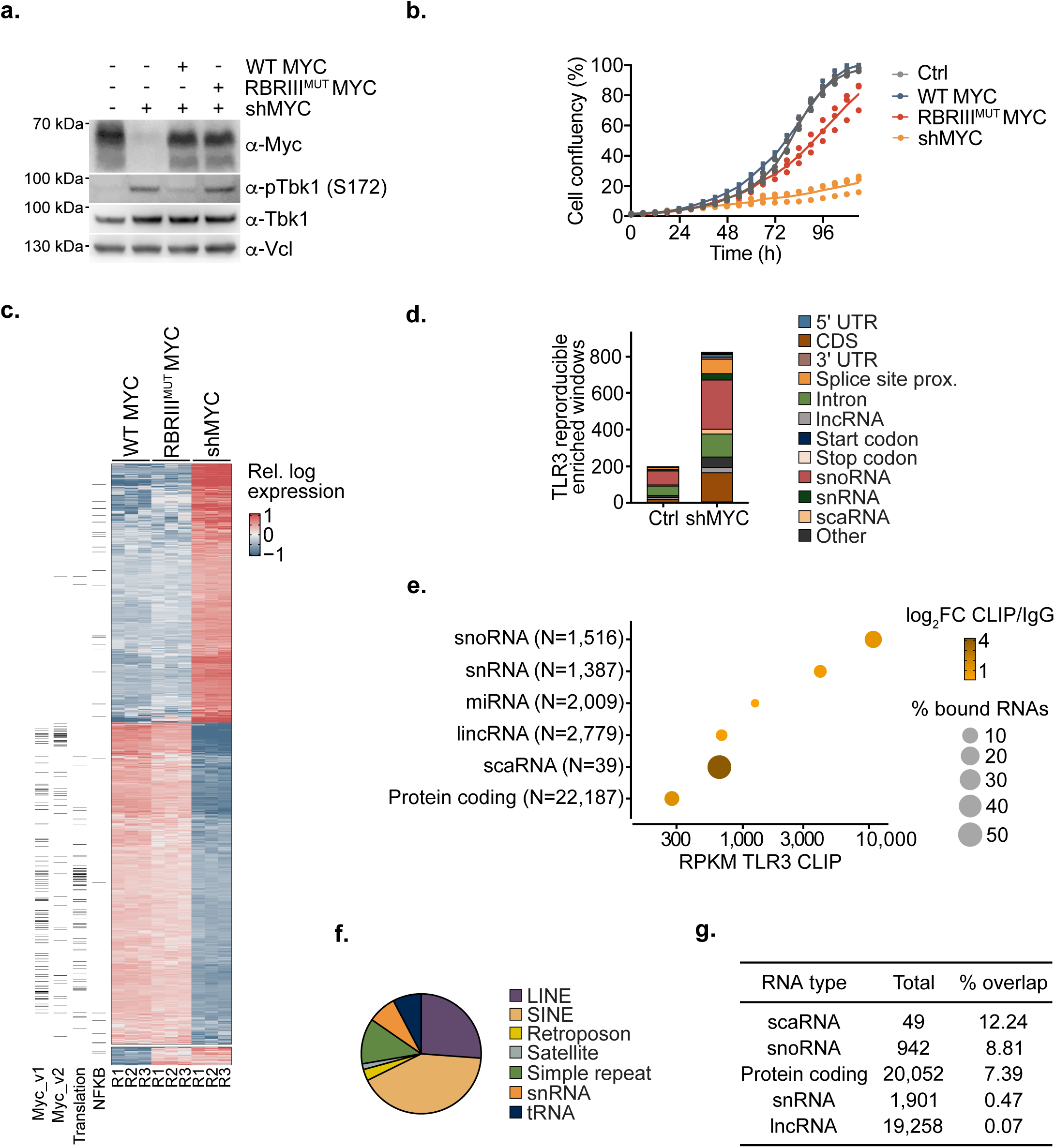
MYC RNA binding suppresses innate immune signaling in PDAC cells. **a.** Immunoblot of KPC^shMYC^ cells stably expressing doxycycline-inducible WT MYC or RBRIII^MUT^ MYC treated with doxycycline (1 µg/ml, 48 h) or EtOH as control (“Ctr” expressing endogenous MYC) (*n*=3). Note that the shMYC targeting MYC is also induced by doxycycline. **b.** Growth curves of KPC^shMYC^ cells stably expressing doxycycline-inducible WT MYC or RBRIII^MUT^ MYC treated with doxycycline (1 µg/mL) or EtOH as control (endogenous MYC) for 5 days. Three biological replicates are shown for each measurement time point (*n*=3). **c.** Heatmap of mRNA sequencing data from KPC^shMYC^ cells after doxycycline (1 µg/ml, 48 h) treatment to induce WT MYC or RBRIII^MUT^ MYC. Exemplary hallmark gene sets and corresponding genes are indicated on the left side of the heatmap (*n*=3). **d.** Number of reproducible enriched windows identified in two TLR3 seCLIP replicates in KPC^shMYC^ cells treated with doxycycline (1 µg/ml, 48 h) or EtOH as control (*n*=2). Number of windows was stratified for the RNA type the windows were located in. **e.** Dot plot showing the reads per kilobase of transcript per million mapped reads (RPKM) of TLR3 seCLIPs per RNA type in doxycycline-treated KPC^shMYC^ cells. The color indicates the log_2_FC of MYC seCLIP signals over IgG cotrols. The size of the dots shows the percentage of transcripts per RNA type enriched in MYC seCLIPs passing a threshold of *p <* 0.05 and log_2_FC CLIP/IgG > 3. The number of RNA molecules in each RNA class was assigned based on the murine annotation (*n*=2). **f.** Pie chart showing the distribution of reproducible repetitive elements on TLR3 upon shMYC induction in KPC^shMYC^ cells identified by Skipper (*n*=2). **g.** Table reporting the percentage of RNAs showing an overlapping binding between MYC and TLR3 as defined of significant enrichment (*p=*0.05) of seCLIPs over IgG controls, and passing a log_2_FC CLIP/IgG threshold of 1. The total number of RNA molecules assigned to each class was based on the human annotation used to analyze the MYC seCLIP conducted in U2OS^MYC-Tet-On^ cells treated with doxycycline (1 µg/mL, 24 h).

In KPC cells, endogenous MYC is also required to suppress the activity of the innate immune sensing kinase, TBK1 (see introduction). In stark contrast to WT MYC, RBRIII^MUT^ MYC was unable to suppress TBK1 autophosphorylation, a hallmark of active TBK1 (Figure 6a). Activation of TBK1 upon depletion of MYC depends on the pattern recognition receptor TLR3, which binds both dsRNA and R-loops^8, 35, 36^. seCLIP experiments showed that TLR3 bound to both coding RNAs and several classes of non-coding RNAs and that the number of enriched RNAs increased for several RNA classes after depletion of MYC (Figure 6d). Normalization of gene length showed showed that small nucleolar RNAs (snoRNAs) were the most strongly bound class of TLR3-bound RNAs (Figure 6e). TLR3 also bound several classes of repetitive elements (Figure 6f), however, due to the large number of individual repetitive elements in each class, it was difficult to quantify the changes for each class of repetitive elements. To determine whether these changes are direct, we surveyed the MYC-bound classes of RNAs and found that in addition to protein-coding RNAs, MYC bound multiple classes of non-coding RNAs, including small nuclear, Cajal-body associated and nucleolar RNAs (Extended Data Figure 6a). Comparison of both datasets showed that a large fraction of RNAs that accumulate on TLR3 after depletion of MYC are directly bound by MYC (Figure 6g and Extended Data Figure 6b, c). Since snoRNAs are the most strongly TLR3-bound class of RNAs (Figure 6e), we focused on them to understand the role of MYC in their turnover.

### MYC prevents loading of misprocessed RNAs onto TLR3

To carry out their function in the nucleolus, snoRNAs assemble with specific proteins to form two distinct classes of snoRNP particles: C/D snoRNPs contain fibrillarin (FBL) and H/ACA snoRNPs contain dyskerin (DKC1)^37^. As expected from their well-characterized localizations in proliferating cells, immunofluorescence imaging showed no overlap in the distribution of MYC and either FBL or DKC1 in unstressed cells (Extended Data Figure 7a). Since FACT is required for transcription by RNA polymerase I, FACT inhibition leads to disintegration of the nucleolus, causing FBL and DKC1 to localize in discrete foci in CBL0137 treated cells (Figure 7a). MYC multimers co-localized with a subset of these foci (Figure 7a).

**Figure 7:**
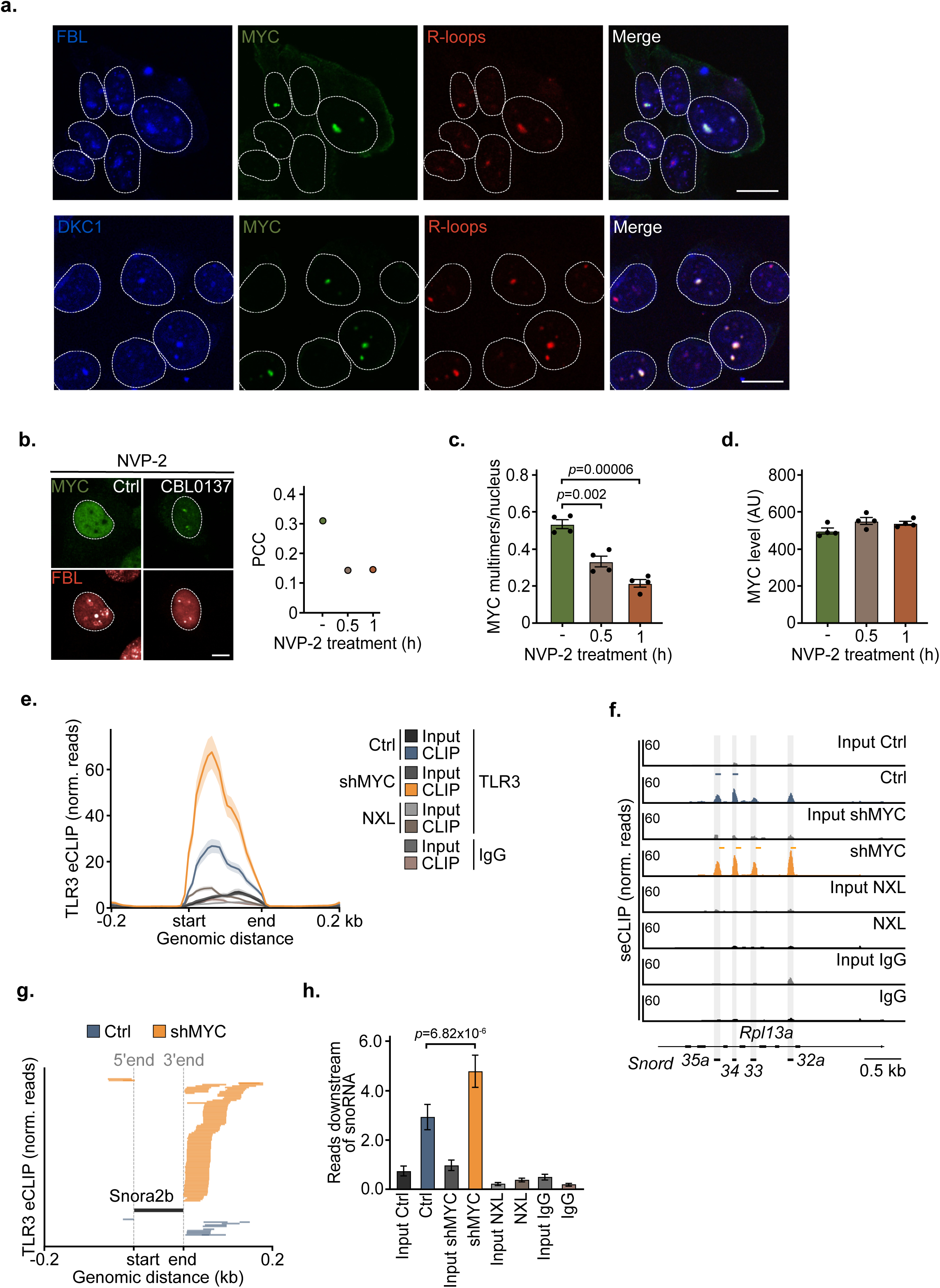
MYC multimers mediate snoRNA processing in stressed cells. **a.** Confocal immunofluorescence of MYC, FBL or DKC1 and R-loops in RNAseH1-HBD-expressing U2OS^MYC-Tet-On^ cells treated with doxycycline (1 µg/mL, 24 h) and CBL0137 (5 µM, 4h) (*n*=4). **b.** Left: Representative immunofluorescence images of MYC and FBL in U2OS^MYC-Tet-On^ cells treated with doxycycline (1 µg/ml, 24 h), NVP-2 (200 nM, 0.5 h or 1 h prior to CBL0137) and CBL0137 (5 µM, 4 h) (*n*=4). Right: The colocalization was quantified applying *PCC*, *t-*test, two-sided (CBL0137 *PCC*=0.31, *p*=2.15×10^-49^, CBL0137; 0.5 h NVP-2 *PCC*=0.14, *p*=6.67×10^-09^; 1 h; NVP-2 *PCC*=0.15, *p*=1.17e^-06^). **c.** Quantification of immunofluorescence showing the number of MYC condensates/nucleus in U2OS^MYC-Tet-On^ cells treated as indicated in b. Shown is the mean of condensates/nucleus ± s.e.m., unpaired, two-sided *t-*test (*n*=4). **d.** MYC nuclear intensity assessed by immunofluorescence of U2OS^MYC-Tet-On^ cells treated as indicated in b and c, presented as mean ± s.e.m. (*n*=4). **e.** Average density plot of TLR3 seCLIPs and inputs at all snoRNAs in KPC^shMYC^ cells treated with doxycycline (1 µg/mL, 72 h) or EtOH as control. IgG seCLIPs and TLR3 eCLIPs from non-crosslinked KPC^shMYC^ cells (NXL) were used as control (*n*=2). **f.** Exemplary browser picture of TLR3 seCLIPs at four snoRNAs located in the *Rpl13a* gene. Samples and treatment conditions as in f. (*n*=2). **g.** Exemplary representation of sequencing reads distributed upstream or downstream of the *Snora2b* gene in KPC^shMYC^ cells treated with doxycycline (1 µg/mL, 72 h) or EtOH as control. Each bar indicates an individual TLR3 seCLIP sequencing read (*n*=2). **h.** Quantification of the number of reads counted within 5 nt downstream of all expressed snoRNAs under the same conditions as in f. Shown are the mean read counts ± s.e.m. Mann-Whitney U, two-sided test was applied.

MYC-positive DKC1 and FBL foci were characterized by the presence of R-loops while MYC-negative foci were devoid of R-loops (Figure 7a). Since MYC proteins are not constituents of mature snoRNP particles, we speculated that MYC multimers associate transiently with snoRNPs during their biogenesis. To test this hypothesis, we made use of the observation that snoRNA host genes are transcribed by RNAPII, which depends on the CDK9 kinase for release from its promoter-proximal pause site and productive elongation^38^. Pre-incubation with a specific CDK9 inhibitor, NVP-2^39^, had no effect on the overall number of snoRNPs containing foci, but rapidly decreased the number of MYC multimers and their co-localization with fibrillarin in CBL0137 treated cells, arguing that MYC associates with snoRNPs during or shortly after snoRNA synthesis (Figure 7b-d). The co-localization of MYC multimers with snoRNPs was rapidly reversible upon removal of CBL0137 (Extended Data Figure 7b), arguing that MYC reversibly associates with nascent snoRNP particles upon FACT inhibition. Both average density plots of all TLR3-bound snoRNAs (Figure 7e) and visual inspection of individual snoRNAs (Figure 7f) confirmed a strong increase in TLR3-binding after depletion of MYC, arguing that a fraction of snoRNAs is exported from the nucleus after MYC depletion. Since RNA binding of MYC promotes the recruitment of the nuclear exosome to R-loops, we surveyed the snoRNAs that accumulate in KPC cells in the absence of MYC for indications of defective exosome function. Indeed, inspection of the ends of TLR3-bound snoRNAs revealed that snoRNAs recovered from MYC-depleted cells often had aberrantly extended 3’-ends, with some snoRNAs carrying significant extensions (Figure 7g,h). We concluded that MYC suppresses the accumulation of snoRNAs and their loading on TLR3 to prevent TBK1 activation and that failure to process or degrade aberrant snoRNAs by the exosome is likely to contribute to their accumulation in the absence of MYC. A model summarizing our findings is shown in Figure 8.

**Figure 8:**
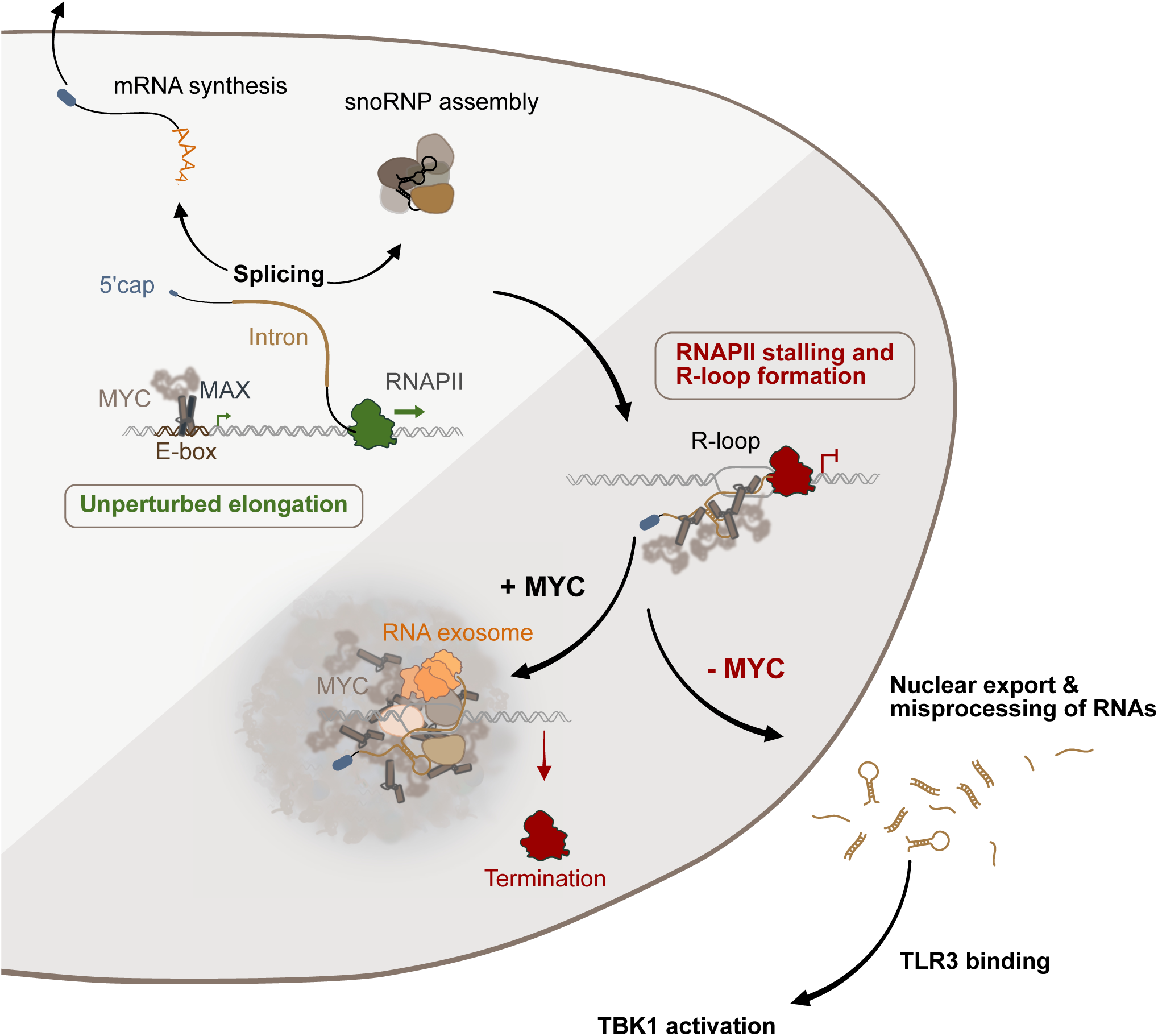
Model summarizing our findings. While MYC binds to active promoters under unstressed conditions, perturbation of transcription elongation leads to relocalization of MYC to RNA structures associated with transcriptional stress such as R-loops or dsRNA. Direct RNA binding of MYC promotes the formation of MYC multimers at these loci, which compartimentalize RNA processing and degradation factors, including the nuclear exosome, thereby preventing the accumulation of immunogenic RNAs.

## Discussion

In response to perturbed transcription elongation, MYC proteins undergo a phase transition and exist in multimeric states with condensate-like properties^14, 40, 41^. We show here that MYC is a direct RNA binding protein and that this phase transition is driven by the accumulation of intronic RNA. Mechanistically, therefore, the phase transition of MYC is similar to other RNA-protein interactions that compartmentalize distinct processes in RNA metabolism^42^ and MYC multimerization likely involves the binding of multiple MYC molecules to adjacent or the same RNA molecule. Importantly, MYC assumes new functionalities upon multimerization. MYC interacts with multiple RNA binding proteins and, upon multimerization, concentrate RNA degrading complexes, in particular the nuclear exosome, a 3’-5’ RNA exonuclease complex, and its NEXT and PAXT targeting complexes around R-loops and potentially other aberrant RNA structures. Collectively, our data argue that MYC multimers compartimentalize stress-induced RNA degradation processes.

A mutant MYC allele that is defective in RNA binding has unaltered transcriptional properties but fails to suppress innate immune signaling. This is because several classes of non-coding RNAs accumulate on TLR3, which recognizes both dsRNA and R-loops, upon MYC depletion. Among these are RNAs derived from repetitive elements as well as snoRNAs and snRNAs. Relative to mature snoRNAs, a fraction of snoRNAs that accumulate on TLR3 upon MYC depletion carry aberrantly elongated 3’-ends, providing direct evidence that they are incompletely processed by the nuclear exosome in the absence of MYC. Mature snoRNAs form dsRNA structures that assemble into snoRNP particles that promote several steps of ribosomal biogenesis. For example, the H/ACA class of snoRNAs specifies uridines in target RNAs for conversion to pseudouridines and modifies ribosomal RNA^43^. MYC proteins associate transiently with nascent snoRNP particles in response to transcription stress.

Several arguments make it unlikely that RNA-bound MYC is involved in physiological snoRNP assembly. First, immunofluorescence of MYC and snoRNP complexes shows no overlap in unstressed, proliferating cells. Second, the assembly pathway of snoRNPs is well understood and is conserved between humans to yeast, which do not have MYC genes^37^. Third, while the exosome is involved in processing the snoRNA host introns, the final nucleotides at the 3’-end of H/ACA snoRNAs are removed by a dedicated nuclease, PARN^44^. We hypothesize, therefore, that the complexes of MYC multimers with the exosome occur when intron processing of the snoRNA host gene is perturbed and contribute to the disassembly of misassembled or misprocessed snoRNP complexes in transcriptionally stressed cells. For example, since the host genes of intron-localized snoRNAs are transcribed into capped mRNAs, MYC-dependent disassembly may prevent the export of aberrant snoRNAs as part of their host mRNAs from the nucleus. Alternatively, incompletely processed snoRNAs may remain part of R-loops, which can be cleaved from the genome and exported from the nucleus to activate TLR3^36^.

In U2OS cells, a fraction of MYC is bound to RNA in unstressed proliferating cells. Similarly, depletion of MYC in PDAC cells enhances the accumulation of 3’-extended snoRNAs in the absence of exogenous stress. Both observations suggest that tumor cells experience endogenous stresses, such as transcription-replication conflicts, that perturb transcription elongation. MYC mutants that are unable to bind RNA and multimerize retain their mitogenic potential and have an unaltered ability to regulate transcription of downstream genes, arguing that they will retain many physiological functions of MYC in cell growth and proliferation. It is likely that the levels of endogenous stress are higher in tumor cells that often proliferate under suboptimal conditions, e.g. with low nucleotide levels. We hypothesize, therefore, that targeting the ability of MYC to bind RNA and multimerize will disrupt immune evasive functions of MYC in tumor development and may therefore open a wide therapeutic window. While the structure of proteins in condensates is likely to be flexible, RNA binding domains are folded and hence targeting this domain and its interaction with RNA using small molecules is feasible and, if successful, enables a specific approach to targeting MYĆs oncogenic functions.

## Extended Data Figure Legends

**Extended Data Figure 1:**
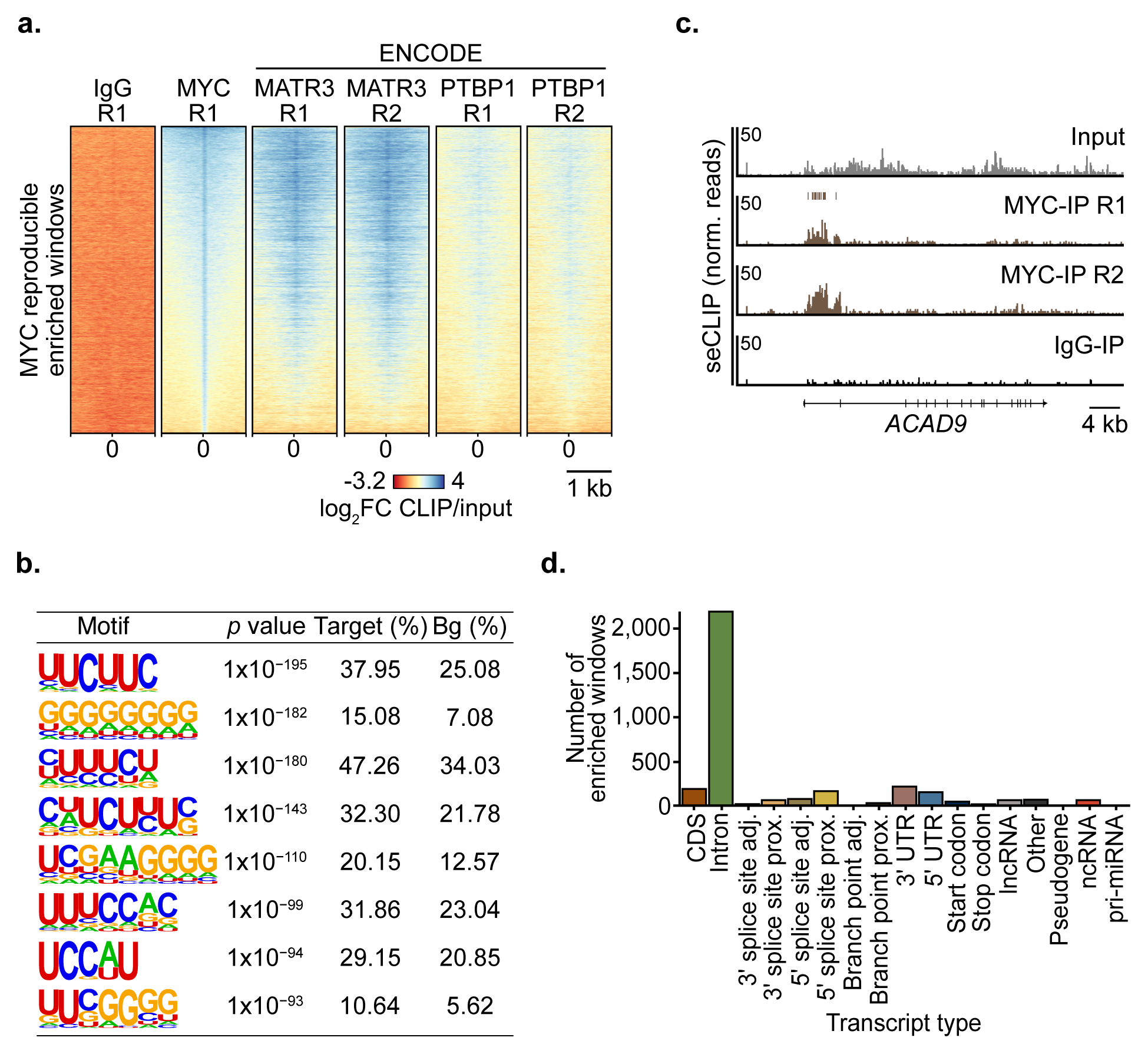
Characterization of MYC RNA binding. **a.** Heatmap of log_2_FC CLIP/input of MYC seCLIPs from K562 cells overlaid with the log_2_FC CLIP/input of publicly available MATR3 and PTBP1 eCLIP data from ENCODE. One of two MYC and IgG seCLIP heatmaps are shown. The MYCR1 graph is the same as in Figure 1 d. **b.** RNA sequence motifs identified by a *de novo* search algorithm (HOMER) at reproducible enriched windows identified in two MYC seCLIP replicates from K562 cells (*n=*2). **c.** Read distribution of MYC seCLIPs in control (EtOH)-treated U2OS^MYC-Tet-On^ cells. IgG seCLIP shown as control (*n*=2). **d.** Number of reproducible enriched windows identified in MYC seCLIPs stratified for different classes of RNA molecules. The experiment was performed in K562 cells (*n=*2).

**Extended Data Figure 2:**
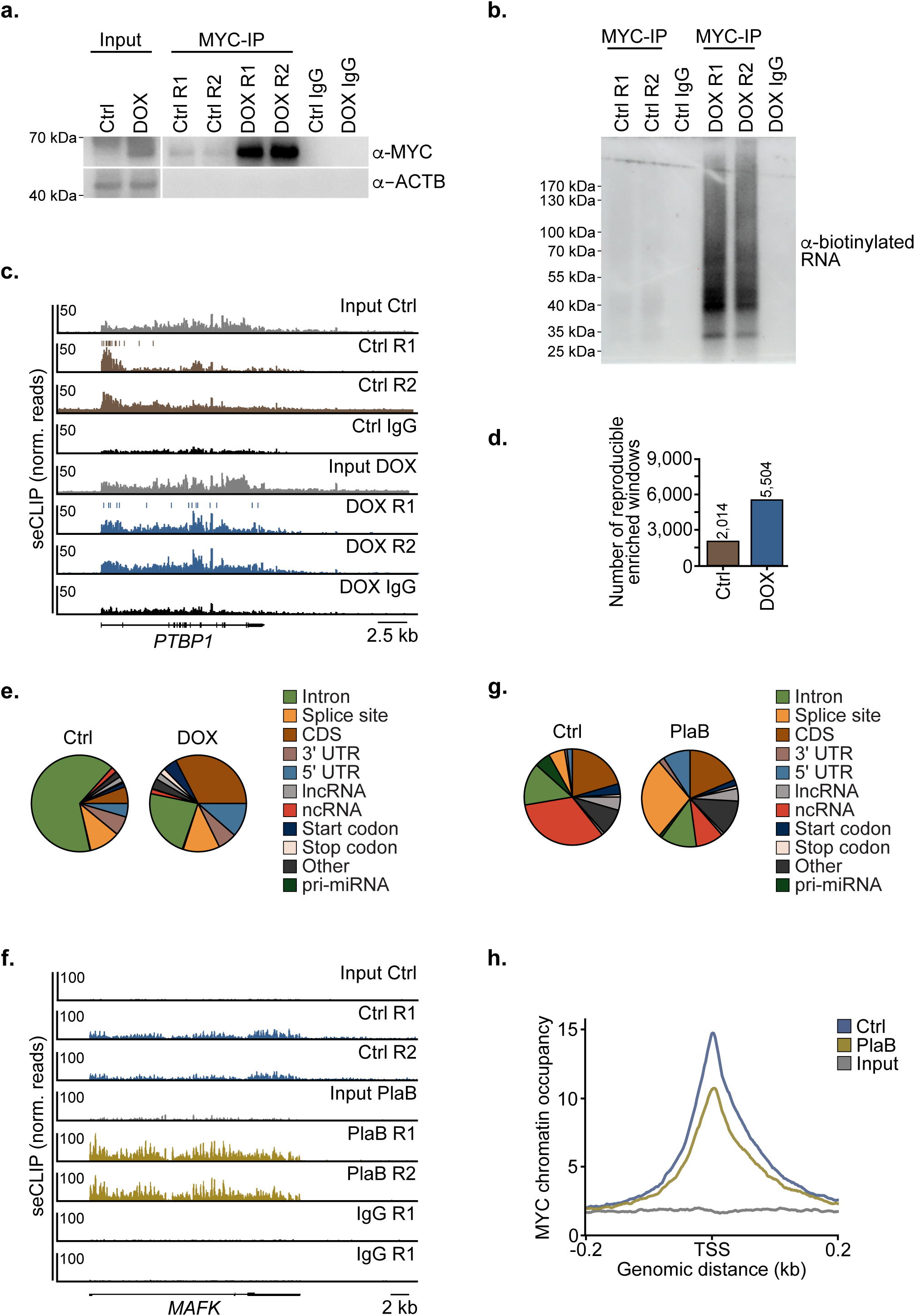
Additional characterization of MYC RNA binding. **a.** Immunoblot of MYC-IPs from U2OS^MYC-Tet-On^ cells treated with doxycycline (1 µg/mL, 24 h) or EtOH as control. Beta-actin (ACTB) served as loading control. Beads coupled to IgG were used to assess non-specific binding. Shown are 2 biological pulldown replicates and one merged input and one IgG control per condition (*n*=2). **b.** Visualization of biotinylated RNA of CLIP experiment MYC pulldowns using HRP-Streptavidin. Cells treated as in a (*n*=2). **c.** Exemplary browser picture of MYC seCLIPs conducted in U2OS^MYC-Tet-On^ cells treated with doxycycline (1 µg/mL, 24 h) or EtOH as control. Bars above Ctrl R1 and DOX R1 MYC seCLIPs show reproducible enriched windows identified in this region (*n*=2). **d.** Number of reproducible enriched windows identified in MYC seCLIPs using Skipper. Treatment conditions as in c. **e.** RNA feature distribution of reproducible enriched windows identified in the different treatment conditions described in c. **f.** Exemplary browser pictures of MYC seCLIPs at the *MAFK* gene conducted in U2OS^MYC-Tet–On^ cells treated with with doxycycline (1 µg/mL, 24 h) and Pladienolide B (PlaB; 1 µM, 4 h) or DMSO as control (*n*=2). **g.** Pie charts showing the RNA feature distribution of reproducible enriched windows in MYC seCLIP experiments described in f. **h.** Average density plot showing MYC chromatin occupancy at the transcription start site (TSS) of 17,086 expressed genes assessed by ChIP-Rx sequencing in U2OS^MYC-Tet-On^ cells treated with doxycycline (1 µg/mL, 24 h) and PlaB (1 µM, 4 h) or DMSO as control (*n*=2).

**Extended Data Figure 3:**
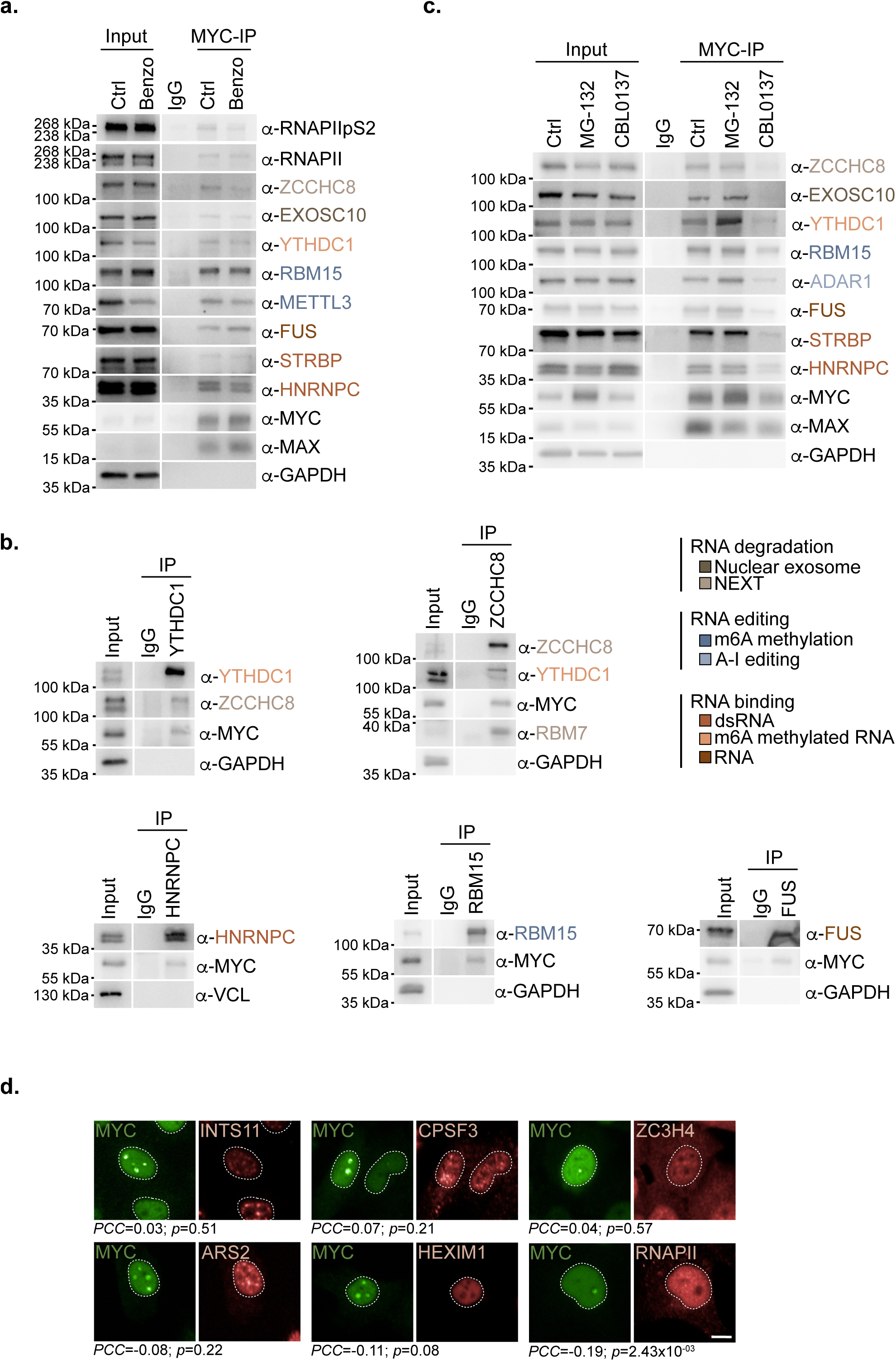
Additional analysis of MYC protein interactions. **a.** Immunoblot of MYC-IPs in U2OS^MYC-Tet-On^ treated with doxycycline (1µg/mL, 24 h) and ± benzonase (“Benzo”, 50 U/ml for 1 h) before precipitation. RNA processing factors were categorizied into canonical RNA binding modes or complexes as indicated by the colour code. GAPDH was used as loading control (*n*=2). **b.** Immunoblots of YTHDC1-, ZCCHC8-, RBM15-, HNRNPC- and FUS-IPs in U2OS^MYC-Tet-On^ cells treated with doxycycline (1µg/mL, 24 h). Samples were treated with RNase inhibitor (50 U/ml) before precipitation. Categorization of factors was performed as described in a. Presented are immunoblots of MYC and known interactors of YTHDC1 and ZCCHC8. GAPDH was used as loading control (*n*=3 for YTHDC1 and ZCCHC8, FUS and HNRNPC *n*=2, RBM15 *n*=1). **c.** Immunoblot of MYC-IPs in U2OS^MYC-Tet-On^ treated with doxycycline (1 µg/mL, 24 h), ± MG-132 (20 µM, 3 h), ± CBL0137 (5 µM, 2.5 h). DMSO served as Ctrl (control). Samples were treated with RNase inhibitor (50 U/mL) before precipitation. Categorization of factors was performed as described in a. GAPDH was used as loading control. (*n*=3 except RBM15, ZCCHC8 *n*=2, ADAR1 *n*=1). **d.** Immunofluorescence of MYC, INTS11, ARS2, CPSF3, Hexim1, ZC3H4 and RNAPII in U2OS^MYC-Tet-On^ cells treated with doxycycline (1 µg/mL, 24 h) and CBL0137 (5 µM, 4 h). The co-localization was quantified using *PCC*, *t-*test, two-sided, Scale bar: 10 µm (*n*=3).

**Extended Data Figure 4:**
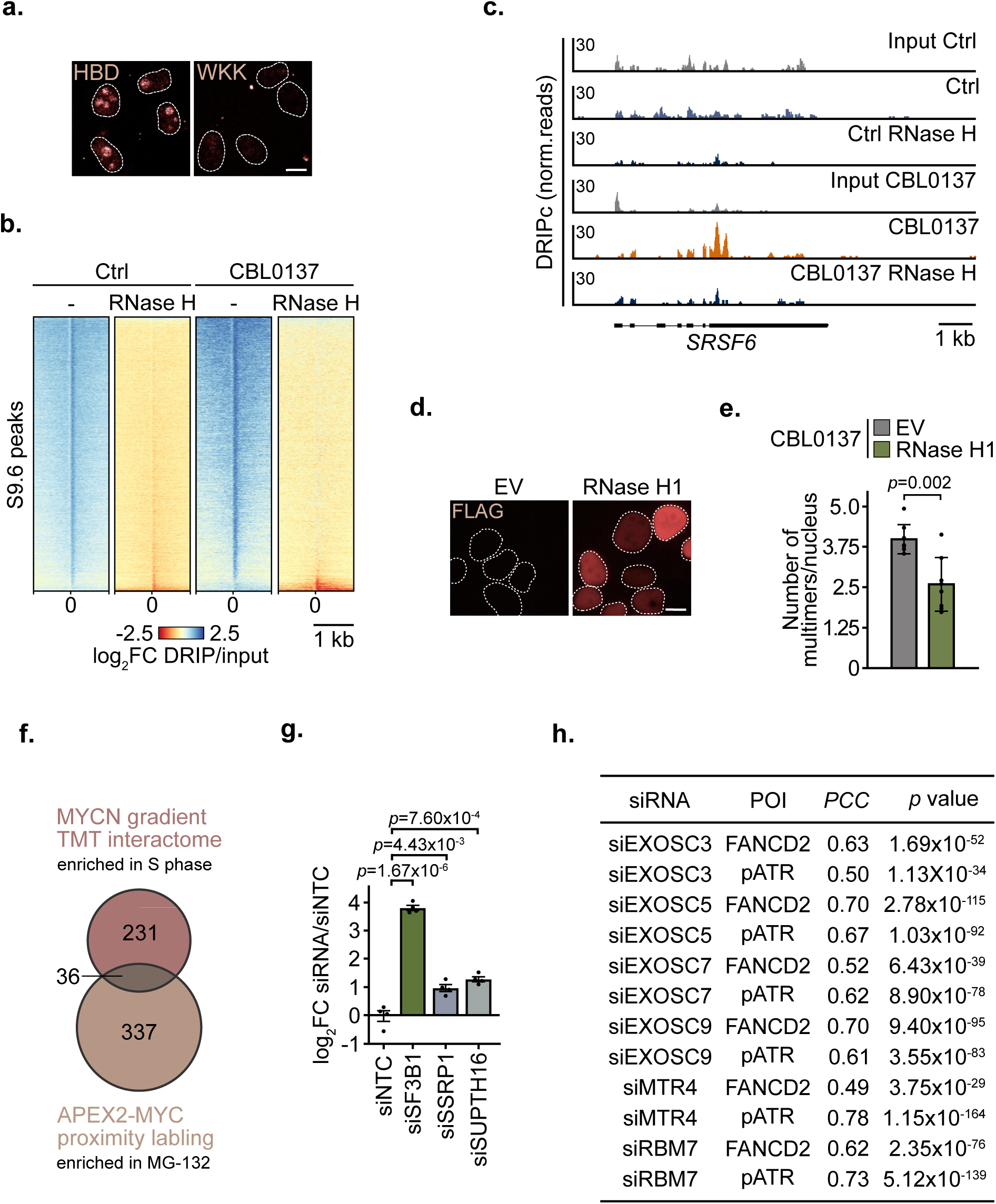
Analyses of R-loops in the context of MYC multimers. **a.** Immunofluorescence of R-loops in RNASeH1-HBD or RNASeH1-WKK-expressing U2OS^MYC-Tet-On^ cells treated with doxycycline (1 µg/mL, 24 h). Scale bar: 10 µm (*n*=3). **b.** Heatmaps showing the log_2_FC DRIP/input of spike-normalized DRIPc sequencing of U2OS^MYC-Tet-On^ treated with doxycycline (1 µg/mL, 24 h) and CBL0137 (5 µM, 2 h) or DMSO as control. Log_2_FC of DRIP/input or DRIP+RNase H/input was plotted at peaks called in Ctrl- and CBL0137-treated condition without RNase H (*n*=1) **c.** Exemplary browser pictures of spike-normalized DRIPc sequencing and input of U2OS^MYC-Tet-On^ treated with doxycycline (1 µg/mL, 24 h) and CBL0137 (5 µM, 2 h) or DMSO as control at the *SRSF6* gene. Samples treated with RNase H were used as control (*n*=1). **d.** Immunofluorescence of a nuclear FLAG-tagged RNase H1 in RNase H1- or EV-expressing U2OS^MYC-Tet-On^ cells treated with doxycycline (1 µg/mL, 24 h). Scale bar: 10 µm (*n*=5). **e.** Bar plot showing the number of MYC multimers per nucleus in control (EV) or NLS-RNase H1-FLAG expressing U2OS^MYC-Tet-On^ cells treated with doxycycline (1 µg/mL, 24 h) and CBL0137 (5 µM, 4 h). Shown is the mean number of condensates/nucleus ± s.d. unpaired, two-sided *t-*test (*n*=8). **f.** Venn diagram based of APEX2 MYC proximity labeling experiment and MYCN gradient TMT interactome. The siRNA library generated for the siRNA screen shown in Figure 4a was based on these datasets and included the indicated overlap of 36 factors as well as 48 factors present in one of the datasets. **g.** Quantification of immunofluorescence showing MYC multimers per nucleus in U2OS^MYC-Tet-On^ treated with siRNAs (48 h) and doxycycline (1 µg/mL, 24 h). Means of biological replicates ± s.e.m. are shown, unpaired, two-sided *t*-test (n=4). **h.** Table showing *PCC* between the partitioning ratio of MYC and FANCD2 or pATR (T1989) upon indicated siRNA treatment (48 h). The *p*-value is the significance level of Pearson’s product-moment correlation *t*-test, two-sided (*n*=3).

**Extended Data Figure 5:**
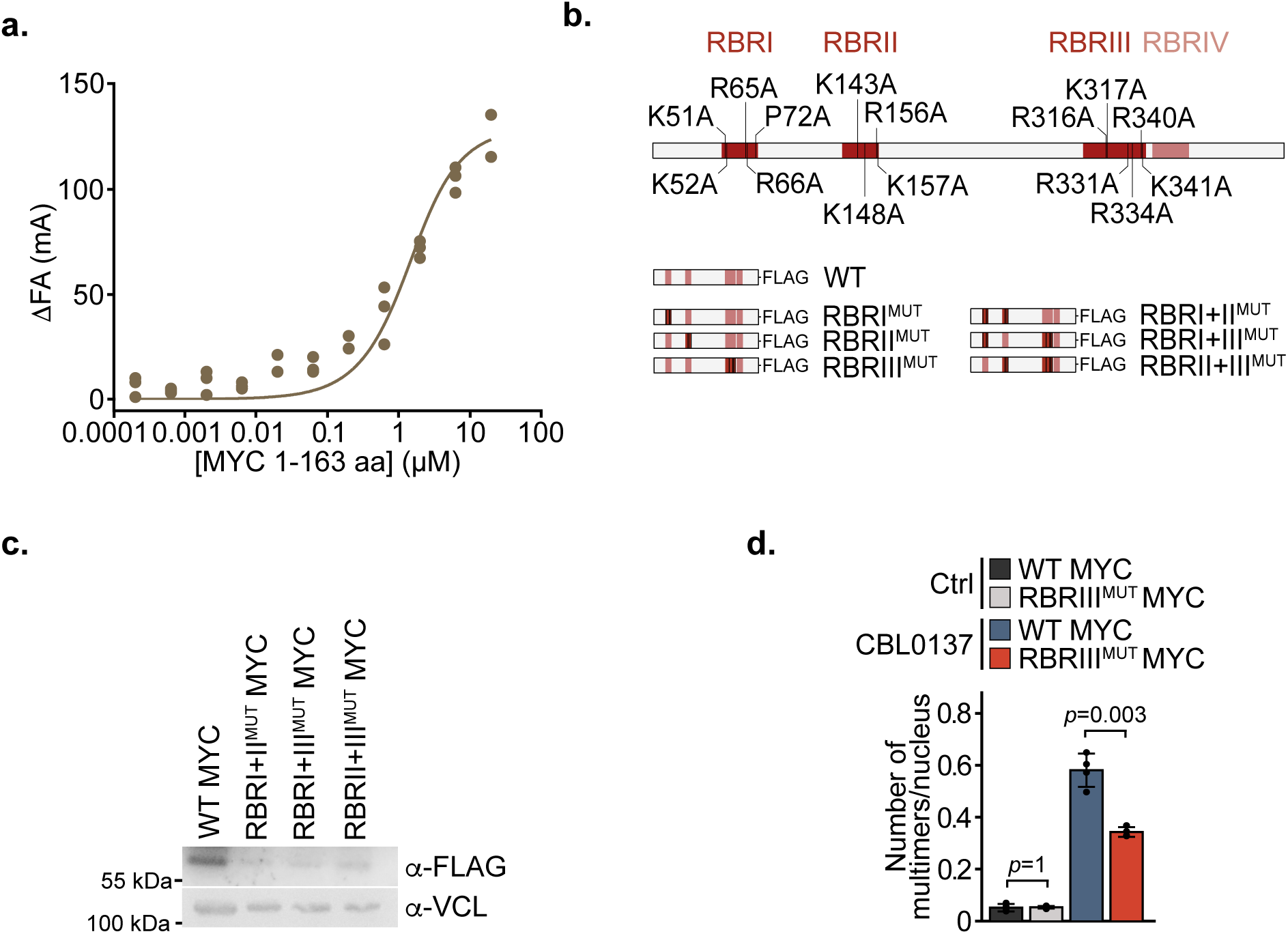
Characterization of RNA binding regions of MYC. **a.** Change in fluorescence anisotropy (ΔFA) relative to 5 nM FITC-labeled RNA using half-log dilutions from 20 μM to 0.2 nM of MYC (1-163 aa). Three biological replicates are shown for each dilution (*n*=3). **b.** Scheme showing the location of mutations introduced (top) to generate different FLAG-tagged MYC RBR mutants (bottom). **c.** Immunoblot of U2OS^MYC-AID^ cells stably expressing doxycycline-inducible FLAG-tagged WT MYC or indicated mutants. Cells were treated with doxycycline (1 µg/mL, 24 h) and IAA (500 µM, 24 h). Vinculin (VCL) is shown as loading control (*n*=2). **d.** Bar graph of immunofluorescence quantifying the number of MYC multimers per nucleus in WT MYC or RBRIII^MUT^ MYC U2OS^MYC-Tet-On^ cells treated with doxycycline (1 µg/mL, 24 h) and CBL0137 (5 µM, 4 h) or DMSO (Ctrl). Presented is the mean number of multimers/nucleus ± s.d. unpaired, two-sided *t-*test (*n*=4).

**Extended Data Figure 6:**
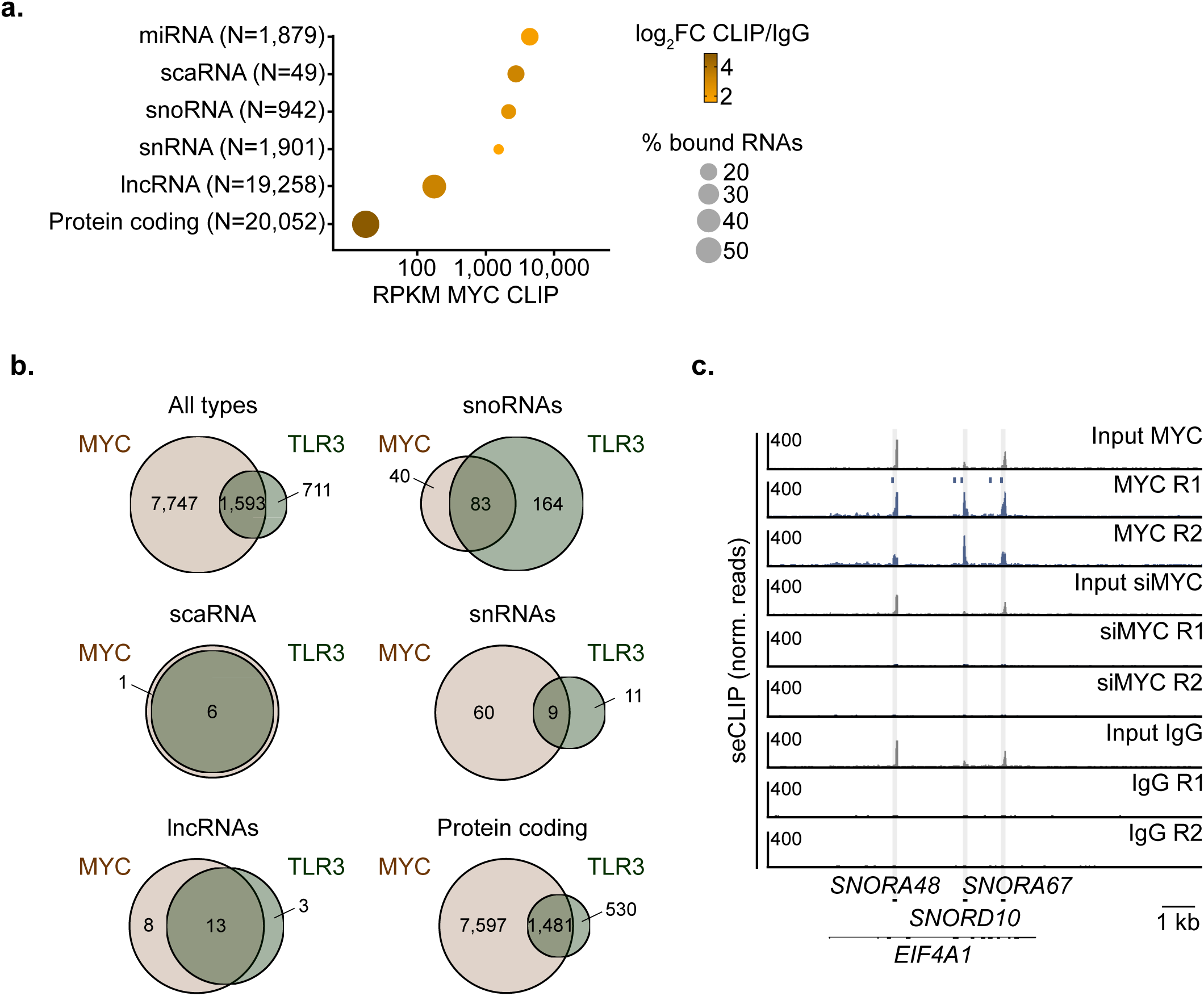
Overlapping binding of TLR3 and MYC to RNA. **a.** Dot plot showing the RPKM of MYC seCLIPs per RNA type in U2OS^MYC-Tet-On^ cells treated with doxycycline (1 µg/mL, 24 h). The color indicates the log_2_FC of MYC seCLIP signals over IgG controls. The size of the dots shows the percentage of transcripts per RNA type enriched in MYC seCLIPs passing a threshold of *p <* 0.05 and log_2_FC CLIP/IgG > 3. The total number of RNAs assigned to each type was determined based on the human annotation (*n*=2). **b.** Venn diagrams showing the overlap between MYC and TLR3 binding to several classes of RNAs as defined as significant enrichment (*p=*0.05) of seCLIPs over IgG controls and passing a log_2_FC CLIP/IgG threshold of 1. The total number of RNA molecules assigned to each class was defined based on the human annotation and used to analyze the MYC seCLIP conducted in U2OS^MYC-Tet-On^ cells, treated with doxycycline (1 µg/mL, 24 h). **c.** Exemplary browser picture of MYC seCLIPs in U2OS^MYC-Tet-On^ cells treated with doxycycline (1 µg/mL, 24 h) at three snoRNA genes located within the *EIF4A1* gene. Bars above the MYC R1 CLIP replicate indicate the position of reproducible enriched windows identified using Skipper (*n*=4).

**Extended Data Figure 7:**
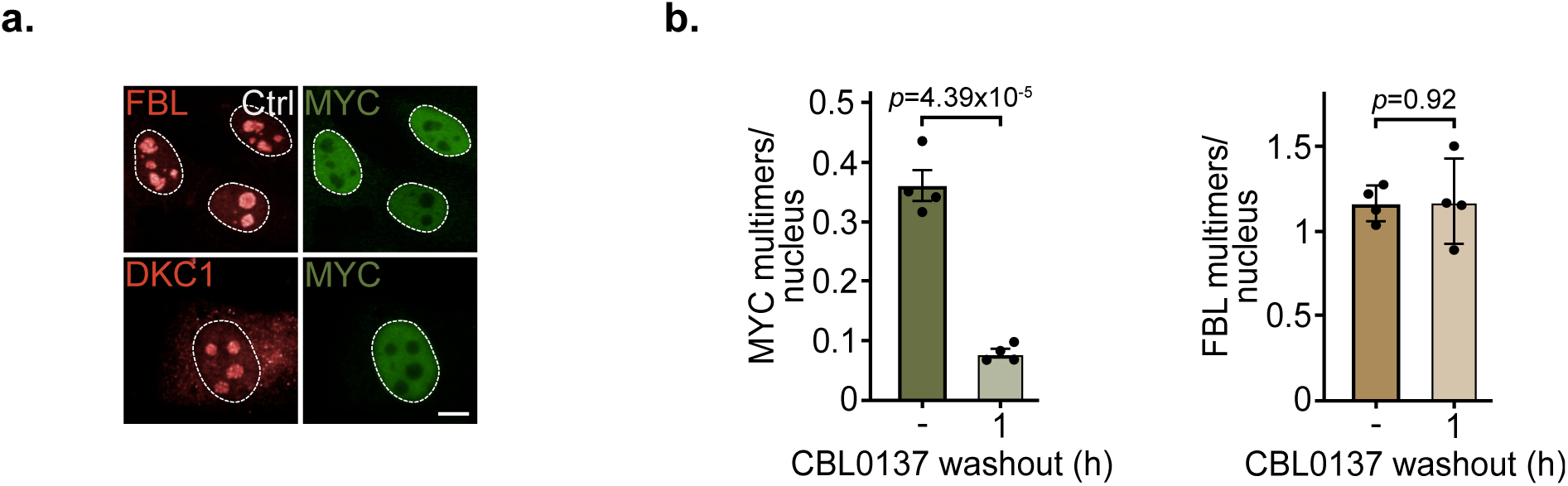
Relationship of MYC multimers to snoRNPs. **a.** Representative immunofluorescence images of MYC, FBL and DKC1 in U2OS^MYC-Tet-On^ cells treated with doxycycline (1 µg/mL, 24 h). Scale bar: 10 µm (*n*=3). **b.** Bar plot quantifying MYC and FBL multimers/nucleus in U2OS^MYC-Tet-On^ cells treated with doxycycline (1 µg/mL, 24 h) and CBL0137 (4 h, 5 µM). CBL0137 washout was performed for 1 h. Shown are mean ± s.e.m. of four biological replicates, unpaired, two-sided *t*-test (*n*=4).

## Methods

### Cell culture and chemical treatment

U2OS, KPC and NIH3T3 cells were grown in DMEM medium containing 10% fetal calf serum (FCS) and 1% penicillin/streptomycin. K562 cells were grown in RPMI-1640 supplemented with 10% FCS and 1% penicillin/streptomycin. *Drosophila* S2 cells were grown in Schneider’s *Drosophila* medium supplemented with 10% FCS and 1% penicillin/streptomycin, and grown at 25 °C. The identity of used cell lines was verified by STR profiling according to the human cell line authentication standard published by ANSI. All cell lines were routinely verified to be *Mycoplasma*-free. Unless stated otherwise, cells were treated with the following reagents at the given concentrations: doxycycline: 1 μg/ml; MG-132: 20 μM; PlaB: 1 µM; CBL0137: 5 µM; NVP-2: 200 nM; IAA: 500 µM.

### Cell growth assays

Cells were seeded in 96-well plates (PerkinElmer) and treated for 5 days with doxycycline (1 µg/ml) or EtOH as control. Medium was refreshed after 72 h. Phase-contrast confluence was monitored by time-lapse microscopy using the Incucyte SX5 system (Sartorius), with 5 fields analyzed every 6 h using 10x magnification The basic analyzer tool in the Incucyte live-cell Imaging software (v2021C) was used to quantify phase object metrics in real-time. The obtained cell confluence values (mean ± s.d.) were visualized with growth curves as percentage over time.

### Transfection and lentiviral infection

siRNA was transfected using RNAiMAX reagent (Thermo Fisher Scientific) according to the manufacturer’s protocol, and cells were harvested after 48 h. Were indicated, doxycycline was added after 24 h of transfection. For siRNA screens shown in Figure 4a and c, wells comprising less than 100 cells were discarded.

For lentiviral infection, U2OS and KPC cells were infected with lentiviral supernatants in the presence of 4 μg/ml polybrene for 24 h. After selection with Neomycin (0.5 mg/ml) for 5 days, cells were seeded for the experiment.

### Protein isolation

Cells were resuspended in HEPES lysis buffer (20 mM HEPES pH 7.9, 200 mM NaCl, 0.5 mM EDTA, 10% glycerol, 0.2% NP-40) to isolate total native proteins. After sonication using four 5-second pulses at 20% power with 10-second pauses, 50 U/ml benzonase or murine RNA inhibitor (NEB) was added to the lysates. Samples were incubated for 1 h at 4 °C on a rotating wheel before clearance by centrifugation. Cells were lysed in RIPA lysis buffer (50 mM HEPES pH 7.9, 140 mM NaCl, 1 mM EDTA, 0.1% v/v SDS, 1% v/v Triton X-100, 0.1% w/v sodium deoxycholate), incubated on ice for 30 min and cleared by centrifugation. A cocktail of phosphatase and protease inhibitors (1:1000; Sigma-Aldrich) was added to HEPES and RIPA lysis buffers. Protein concentrations were assessed by the BCA or Bradford assay.

### Co-IPs and immunoblotting

For co-immunoprecipitation, 20 μl Protein A/G beads were used in a 1:1 ratio per IP. After washing three times with 5 mg/ml BSA/PBS, the beads were coupled with 2.5 μg of antibody against MYC (abcam, ab32072), ZCCHC8 (Thermo Fisher Scientific, A301-806A), HNRNPC (Proteintech, 11760-1-AP), RBM15 (Proteintech, 10587-1-AP), FUS (Thermo Fisher Scientific, A300-294A) or YTHDC1 (abcam, ab264375) in 5 mg/ml BSA/PBS solution and incubated overnight at 4 °C on a rotating wheel. IgG was used as a control. Lysates were pre-cleared for 1 h at 4 °C by adding 20 μl Protein A/G beads at a 1:1 ratio per IP. Then, 1-2 mg of lysate was added to the pre-coupled antibody-bead mixture. After 4-6 h, samples were washed four times with HEPES lysis buffer and eluted with 2x Lämmli buffer (15 mM Tris pH 6.8, 3% SDS, 0.015% bromophenol blue, 10% glycerol, 1.5 mM 1,4-dithiothreitol). Samples were heated at 95 °C for 5 min before loading onto Bis-Tris gels and wet transfer to a PVDF membrane (Merck). The membranes were then blocked with 5% BSA in TBS-T for 1 h and incubated with the appropriate primary antibodies overnight at 4 °C. The membranes were incubated with HRP-conjugated secondary antibodies matching the host organism of the primary antibodies for 1 h at RT. Fusion FX (VILBER) was used for image acquisition.

### Immunofluorescence

For high-throughput imaging, cells were seeded in 96-well plates (Revvity). For immunofluorescence of R-loops, cells were pre-extracted with CSK buffer (10 mM Pipes (pH 7.0), 100 mM NaCl, 300 mM sucrose, 3 mM MgCl_2_, 0.7% Triton X-100) for 3 min at RT prior to fixation. For all other stainings, cells were directly fixed with 4% paraformaldehyde/PBS for 10 min at RT, permeabilized with 0.1% Triton-X100/PBS for 10 min at RT, and blocked with 5% BSA/PBS for 1 h at RT. Primary antibodies in 5% BSA were then added and incubated overnight at 4 °C. The cells were subsequently washed with PBS and secondary antibodies corresponding to the host of the primary antibody were added. After incubation for 1 h at RT, cells were counterstained with Hoechst33342 (Sigma-Aldrich). Cells were kept in PBS at 4 °C until images were captured with the Operetta CLS High-Content Imaging System at 40x and 60x magnification using a water immersion objective. Image processing was performed with the Harmony High Content Imaging and Analysis Software (Revvity).

For confocal microscopy cells were seeded on 8-well □-slides (ibidi). Immunofluorescence was conducted as described before and cells were covered with mounting medium (ibidi). Slides were scanned utilizing the Leica SP2 inverted confocal laser scanning microscope equipped with diode, argon, and helium neon lasers (405 nm, 458 nm, 476 nm, 488 nm, 496 nm, 514 nm, 561 nm, 633 nm) using an HCX PL APO Leica 63x immersion oil objective (NA 1.4).

### Proximity Ligation Assay (PLA)

Cells were grown in 384-well plates (Revvity) and treated as indicated prior to fixation with 4% paraformaldehyde/PBS for 10 min at RT, permeabilization with 0.1% Triton-X100/PBS for 10 min at RT, and blocking with 5% BSA/PBS for 1 h at RT. Primary antibodies in 5% BSA were then added and incubated overnight at 4 °C. The next day, PLA was performed using the Duolink In Situ Kit (Sigma-Aldrich) according to the manufacturer’s instructions. In brief, cells were washed with TBS-T and incubated with plus and minus PLA probes specific for rabbit and mouse antibodies for 1 h at 37 °C. The cells were subsequently washed with Wash Buffer A (Sigma-Aldrich) and the ligase reaction was performed at 37 °C for 30 min. The *in situ* PCR amplification was then conducted for 2 h at 37 °C, the cells were rinsed with wash buffer B (Sigma-Aldrich), and the nuclei were counterstained with Hoechst33342 (Sigma-Aldrich). Images were acquired with the Operetta CLS High-Content Imaging System at 40x magnification using a water immersion objective. Image processing was done using Harmony High Content Imaging and Analysis Software (Revvity).

### Peptide library synthesis and microarray printing

MYC (MYC_HUMAN, Uniprot P01106) residues (1-454) were synthesized as 25 amino acid (aa) long peptides as an overlapping library. Overlap was 22 aa and off-set 3 aa, resulting in 144 peptides. μSPOT peptide array synthesis was conducted as outlined previously ^16^ ^45^. In brief, parallel and automated (MultiPep Rsi, CEM, USA) solid phase peptide synthesis was conducted on 9-fluorenylmethyloxycarbonyl-β-alanine (FMOC-β-Ala) modified cellulose, laser-cut in 4 mm wide discs. All reagents, chemicals and building blocks were synthesis grade or better (IRIS Biotech GmbH, Germany) and used without any further purification. Loading per disk was ∼130 nmol. FMOC was removed using 20% v/v piperidine in dimethylformamide (DMF) and discs were washed with DMF and ethanol (EtOH). Preactivated FMOC protected aa (0.5 M), coupling agent N,N′-diisopropylcarbodiimide (1 M) and ethyl 2-cyano-2-(hydroxyimino)acetate/Oxymapure (1 M) was dispensed onto discs followed by washing with three times. Unreacted free amine group were capped with a 1:25 dilution of acetic anhydride in DMF and washed with DMF followed by EtOH. After N-terminal FMOC removal discs were washed and dried and transferred to 96-well plates for further processing. Side chain protecting groups were removed with cleavage cocktail containing 90% trifluoracetic acid (TFA), 3% triisoproplysilane (TIPS), 2% dichloromethane and water 5% (all v/v) for 2 h on a rotary shaker at 1000 rpm. The cleavage mix was discarded and a mixture containing TFA 88.5%, trifluoromethanesulfonic acid (TFMSA) 4%, water 5% and TIPS 2.5% (all v/v) was added and incubated overnight rotating at 500 rpm. Peptide cellulose conjugates were precipitated with chilled diethyl ether, centrifuged at 4 °C and washed at least 2 times followed by drying. Conjugates were reconstituted in DMSO, later 2 parts of PCC were mixed with saline-sodium citrate buffer (150 mM NaCl, 15 mM trisodium citrate, pH 7.0) and transferred to 384 well plate and printed using a contact printer (CEM GmbH, USA) as duplicates of 50 nl drops on coated blank slides (76 mm x 26 mm, Intavis Peptide Services GmbH) and left for drying overnight.

### Microarray binding assay

Peptide library arrays were protected from stray light using aluminum foil during assay. Arrays were equilibrated in PBS-T buffer (137 mM NaCl, 2.7 mM KCl, 10 mM Na_2_HPO_4_, 1.8 mM KH_2_PO_4_, pH 7.4, 0.1% v/v Tween20) for 0.5 h. Buffer was removed and replaced by 5’ conjugated Cy5 RNA sequence (UGGGCCUGGAAUAGUGGGGGCCCAGGGGAAGCUGCGAUGUGCUUCUGAGUGUUUGGAACGAG) dissolved in PBS-T buffer. After 15 min, the arrays were washed with PBS-T, the arrays were imaged on Amersham ImageQuant 800 (Cytiva, Danaher, USA) with exposure time between 0.1-10 seconds. Images were analyzed with FIJI with microarray profile addon (OptiNav), average and derivation was derived from 6 fields from duplicates of 3 arrays.

### Expression and Purification of MYC constructs

cMYC constructs were expressed in BL21 (DE3) RIL LOBSTR cells. Overnight cultures were used to inoculate 4L of 2xYT media and cells were grown at 37 °C, shaking in baffled flasks until reaching an OD600 of 0.4-0.6. The temperature was then lowered to 30 °C (cMYC 304-354) or 18 °C (cMYC 1-163) and protein expression was induced by the addition of 0.5 mM IPTG. Cells were then grown for 5 h (cMYC 304-354) or 16 h (cMYC 1-163). Cells were collected by centrifugation and resuspended in lysis buffer (300-500 mM NaCl, 20 mM Tris-HCl pH 7.9 at 25 °C, 10% (v/v) glycerol, 30 mM imidazole pH 8.0, 5 mM beta-mercaptoethanol, 0.284 μg/m leupeptin, 1.37 μg/ml pepstatin A, 0.17 mg/ml PMSF, and 0.33 mg/ml benzamidine). Resuspended cells were either used directly for protein purification or snap frozen in liquid nitrogen and stored at -80 °C.

For protein purification, all steps were performed at 4 °C unless otherwise noted. Cells were lysed by sonication and lysates were clarified by centrifugation. Clarified lysate was applied to a 5 mL HisTrap column (Cytiva) equilibrated in lysis buffer. The column was washed with lysis buffer and additionally washed with a high salt buffer (lysis buffer with 1 M NaCl) for 5 column volume (CV) before returning to lysis buffer. Protein was eluted from the HisTrap column with nickel elution buffer (200-500 mM NaCl, 20 mM Tris-HCl pH 7.9, 500 mM imidazole pH 8.0, 10% glycerol, and 5 mM BME) onto a 10 ml amylose column (New England Biolabs). The amylose column was washed with lysis buffer. Protein was eluted from the amylose column in amylose elution buffer (200-500 mM NaCl, 20 mM Tris-HCl pH 7.9, 30 mM imidazole pH 8.0, 116 mM maltose, 10% glycerol, and 5 mM BME). Protein purity was assessed by SDS-PAGE followed by Coomassie staining. Fractions containing cMYC were pooled and concentrated with an Amicon Millipore ultrafiltration device (10000 MWCO). Concentrated protein was applied to a HiLoad Superdex 200 16/600pg column (GE) equilibrated in size exclusion buffer (200 mM NaCl, 20 mM Tris-HCl pH 7.9, 10% glycerol, and 1 mM DTT). Protein purity was assessed by SDS-PAGE followed by Coomassie staining. Fractions containing cMYC were pooled and concentrated with a 10 MWCO Amicon Millipore ultrafiltration device. Protein concentration was determined using the calculated extinction coefficient and the measured absorbance at 280 nm. Protein was aliquoted, snap frozen in liquid nitrogen, and stored at - 80 °C until use.

### Fluorescence anisotropy (FA) RNA binding assay

5′ 6-FAM labelled RNAs were purchased from IDT with the following PTDSS2 oligo sequence: 5’-GUGCAUGGGCAUGUGUGCUGUGUGUGUGCAGGCGACCGUAAACA-3’. Briefly, cMYC was titrated and mixed with 5 nM of fluorescently labelled RNA in a final buffer containing 50 mM NaCl, 20 mM Tris-HCl pH 7.9, 5% (v/v) glycerol, 1 mM DTT. Reactions were incubated at RT in the dark for 20 min. 18 μl of the reaction was transferred to a 384-well plate (Corning) and fluorescence anisotropy was detected using a Tecan Spark plate reader with a gain of 150, excitation of 470 ± 20 nm and emission of 518 ± 20 nm, Z-height of 15000 μm. Fluorescence anisotropy values were normalized by subtracting the anisotropy detected for the RNA alone from each well with protein and RNA. Experiments were performed at least 3 times. Data were graphed in GraphPad Prism (version 9.5.1) and fit using a single-site quadratic binding equation.

### Visualization of crosslinked RNA

Samples for visualization of cross-linked RNA were either prepared in an independent experiment or taken from seCLIP samples which were prepared for sequencing. Cell lysates and pulldowns were performed as described in^19^. Afterwards biotinylated cytidine (bis)phosphate was added to the dephosphorylated RNA samples using a RNA ligase high-concentration enzyme in an on-bead ligation reaction conducted overnight at 16 °C. Beads were subsequently washed once with seCLIP high-salt wash buffer followed by a transition wash and three more washes with seCLIP wash buffer. NuPAGE LDS buffer (Thermo Fisher Scientific) containing DTT (98 mM) was added and incubated at 70 °C for 10 min. After separating the protein-RNA complexes via gel electrophoresis utilizing BisTris gels, samples were transferred to nitrocellulose membranes. The biotinylated RNA was stained using the chemiluminescent nucleic acid detection module kit (Thermo Fisher Scientific) following the manufacturer’s instructions. In brief, membranes were incubated with pre-warmed blocking buffer for 15 min at RT with gentle shaking, followed by a 15 min incubation with the streptavidin-HRP conjugate in blocking buffer. Membranes were then washed four times with 1x washing solution and incubated with substrate equilibration buffer for 5 min. Chemiluminescent substrate working solution was added for 5 min protected from light and images were taken using the Fusion FX.

### Spike-in chromatin immunoprecipitation and sequencing (ChIP-Rx)

To crosslink cells following treatment, 1% v/v formaldehyde was added to the cell culture medium and incubated 5 min rotating at RT. 125 mM glycine were added to stop the reaction and incubated for 5 min rotating at RT. Cells were then harvested in ice-cold PBS containing protease and phosphatase inhibitors (Sigma Aldrich). Cells were centrifuged (1,500 rpm, 20 min, 4 °C) and the pellet was resuspended in ChIP lysis buffer I (5 mM PIPES pH 8.0, 85 mM KCl, 0.5% NP-40). For spike-normalization, 10% murine NIH3T3 cells were added based on the cell number. After 20 min incubation on ice, nuclei were pelleted by centrifugation (1,500 rpm, 20 min, 4 °C) and subsequently lysed by incubation in ChIP lysis buffer II (10 mM Tris pH 7.5, 150 mM NaCl, 1 mM EDTA, 1% NP-40, 1% sodium deoxycholate, 0.1% SDS) for 10 min on ice. The chromatin was fragmented using the Covaris Focused Ultrasonicator M220 (50 min; cycles/burst, 200; peak power, 75; duty factor, 10), and the required fragment size of 150-300 bp was validated by agarose gel electrophoresis. The fragmented chromatin was collected by centrifugation (14,000 rpm, 20 min, 4 °C). For each IP, 100 µl of each Protein A and Protein G-coupled Dynabeads (Thermo Fisher Sientific) were pre-incubated overnight with 15 µg antibody in the presence of 5 mg/ml BSA at 4 °C. Subsequently, beads were added to the chromatin and incubated overnight at 4 °C on a rotating wheel. Immunoprecipitated protein-chromatin complexes were washed 3 times each with ice-cold ChIP wash buffer I (20 mM Tris pH 8.1, 150 mM NaCl, 2 mM EDTA, 1% Triton X-100, 0.1% SDS), ChIP wash buffer II (20 mM Tris pH 8.1, 500 mM NaCl, 2 mM EDTA, 1% Triton X-100, 0.1% SDS), ChIP wash buffer III (10 mM Tris pH 8.1, 250 mM LiCl, 1 mM EDTA, 1% NP-40, 1% sodium deoxycholate; including a 5 min incubation at 4 °C), and once with ice-cold TE buffer. Protein-chromatin complexes were eluted twice by incubating in 150 µl elution buffer 100 mM NaHCO_3_, 1% SDS) for 15 min at RT. To remove potential RNA contaminations, IP and input samples were treated with RNase A for 1 h at 37 °C. Samples were de-crosslinked by incubation at 65 °C shaking overnight, and then treated with Proteinase K for 2 h at 45 °C to degrade the protein. The amount of DNA was quantified using Quant-iT PicoGreen dsDNA assay (Thermo Fisher Scientific) and libraries were generated using the NEBNext Ultra II DNA Library Prep Kit for Illumina (New England Biolabs) following manufacturer’s instructions.

### seCLIP sequencing

seCLIP (single-end enhanced cross-linking and immunoprecipitation) sequencing was performed as described in ^19, 46^. All seCLIP experiments were conducted in duplicates with one pooled size-matched inputs per condition. Per IP sample, 30-40 million cells were crosslinked with UV-C light (254 nm, 2 min, 400 mJ/ cm^2^). Not UV-crosslinked cells were used as negative control. Cells were lysed in seCLIP lysis buffer (50 mM Tris-HCl pH 7.4, 100 mM NaCl, 1% (vol/vol) Igepal CA-630, 0.1% (vol/vol) SDS and 0.5% (wt/vol) sodium deoxycholate) supplemented with a protease inhibitor cocktail (Sigma-Aldrich) and 440 U/ml murine RNase inhibitor (NEB), and 10% of crosslinked murine NIH 3T3 cells were added to each sample as spike-in. Samples were sonicated using the M220 Focused-ultrasonicator (Covaris) for 3 min. Next, samples were treated with 40 U RNase I (Thermo Fisher Scientific) and 10 U Turbo DNase for 5 min at 37 °C shaking at 1200 rpm. After centrifugation at 15,000 g for 10 min at 4 °C, supernatants were added to pre-coupled protein A/G Dynabeads (62.5 µl each per IP) to the antibody of interest (10 µg per sample) and incubated rotating at 4 °C overnight. 2% v/v of the lysate (20 µl) including beads were taken as SMInput, both for the RNA gel and for the diagnostic western blot. The remaining lysate was magnetically separated and washed 2 times with high-salt wash buffer (50 mM Tris-HCl pH 7.4, 1 M NaCl, 1% (vol/vol) Igepal CA-630, 1 mM EDTA, 0.1% (vol/vol) SDS and 0.5% (wt/vol) sodium deoxycholate), followed by a transition wash which was conducted by adding 500 µl of high-salt wash buffer, mixing and then adding 500 µl of wash buffer (20 mM Tris-HCl pH 7.4, 10 mM MgCl_2_, 0.2% (vol/vol) Tween-20 and 5 mM NaCl). Three more washes with wash buffer were conducted, before the immunoprecipitated RNA was dephosphorylated on-beads using the FastAP and PNK enzymes. Therefore, 1X FastAP buffer (Thermo Fisher Scientific), 80 U murine RNase inhibitor (NEB), 4 U Turbo DNase (NEB) and 3 U FastAP enzyme (Thermo Fisher Scientific) were added in a total volume of 50 µl and incubated for 20 min at 37 °C shaking at 1,200 rpm. Without removing the phosphorylation mix, 150 µl PNK master mix (1x PNK7 buffer (700 mM Tris-HCl pH 7, 100 mM MgCl_2_), 40 U PNK4 enzyme) were added and incubated for another 20 min at 37 °C (shaking at 1,200 rpm). Samples were then washed with 200 µl seCLIP high-salt wash buffer once, followed by a transition wash and three further washings with seCLIP wash buffer. 10% of the IP samples were saved for biotin-labeling to visualize the RNA cross-linked to the protein of interest. To the remaining RNA, InvRiL19 RNA adapters (IDT) were ligated by incubating the samples with 25 µl 3’RNA linker mix (1.2x RNA ligase buffer (no DTT), 1.2 µM ATP, 3.6% DMSO, 0.024% v/v Tween-20, 18% w/v PEG 8000, 0.8 U murine RNase inhibitor (NEB), 72 U T4 RNA ligase high-concentration enzyme) rotating at RT for 75 min. Beads were then washed as follows: 1x wash buffer, transition wash to high-salt wash buffer, 1x high-salt wash buffer, transition wash to wash buffer, 3x wash buffer. After the final wash, 20% per IP sample was kept for the diagnostic western blot. To all IP and SMInput samples, NuPAGE LDS buffer (Thermo Fisher Scientific) containing DTT (98 mM, freshly prepared) was added before samples were incubated at 70 °C for 10 min. After magnetically separating the beads from the lysates, RNA-protein complexes were separated by gel electrophoresis using 8% Bis-Tris gels, and transferred onto nitrocellulose or methanol-activated PVDF membranes for the RNA gels and western blots, respectively. The diagnostic western blot was used as a guide to cut out the region of the membrane starting from the observed size of the protein of interest and extending until the maximal expected size of the RNA-protein complexes (but at least ∼75 kDa larger). Membrane pieces were then sliced into ∼2 mm x 2 mm squares and transferred into a fresh tube. To digest the protein, membranes were incubated with 24 U proteinase K (Roth) in PKS buffer for 20 min at 37 °C followed by an incubation at 50 °C for another 20 min. RNA was extracted using the RNA Clean and Concentrator-5 kit (Zymo) following the manufacturer’s instructions, and RNA was eluted in 10 µl H_2_O. Next, SMInput samples were dephosphorylated by incubating each SMInput RNA sample with 20 µl of dephosphorylation master mix (1x FastAP buffer (Thermo Fisher Scientific), 40 U murine RNase inhibitor (NEB), 2 U FastAP enzyme (Thermo Fisher Scientific) incubating at 37 °C for 20 min shaking at 1,200 rpm, followed by another 20 min incubation at 37 °C (1,200 rpm) in PNK master mix (1.05x PNK7 buffer, 5.26 mM DTT, 2 U Turbo DNase (NEB), 40 U T4 PNK enzyme). RNA was purified using the Clean & Concentrator 5 kit (Zymo) as described before. To append the SMInput RNA with 3’ RNA adapters, 1.5 µl 100% DMSO and 0.5 µl InvRil19 adapter were added per sample and incubated at 65 °C for 2 min. 13.5 µl of SMInput RNA ligation master mix (0.976x RNA ligase buffer (NEB), 0.976 mM ATP, 2.93% DMSO, 0.02% v/v Tween-20, 14.63 w/v PEG 8000, 12 U murine RNase inhibitor, 36 U T4 RNA ligase high-concentration enzyme) were added and samples were incubated at RT for 60 min on a rotating wheel. To purify the RNA, 15 µl MyOne Silane Dynabeads per sample were washed ones with RLT buffer (Qiagen), resuspended in RLTW buffer (RLT + 0.025% v/v Tween-20) and added to the RNA. An equal volume of 100% EtOH was added and samples were incubated at RT for 10 min, before samples were washed three times with 80% v/v EtOH. After the final wash, beads were tried and subsequently eluted in 9.5 µl of TT elution buffer (10 mM Tris-HCl pH 7.4, 0.1 mM EDTA, 0.01% (vol/vol) Tween-20). All IP and SMInput samples were next reverse transcribed using the SuperScript III enzyme. For that, 1 µl of 5 µM InvAR17 RT primer and 1 µl of 10 mM dNTPs were added per RNA sample and incubated for 2 min at 65 °C. 10 µl of reverse transcription master mix (1x first strand buffer (Invitrogen), 5 mM DTT, 8 U murine RNase inhibitor (NEB), 120 U Superscript III enzyme (Invitrogen)) were added to each sample and incubated at 55 °C for 20 min. 2.5 µl ExoSAP-IT (Thermo Fisher Scientific) were used to remove unincorporated dNTPs and RT primers (37 °C, 15 min) before 1 µl of 0.5 M EDTA was added. To remove RNA stands, 3 µl of 1 M sodium hydroxide were added and incubated at 70 °C for 10 min. 3 µl of 1 M hydrochloric acid were added to neutralize the sodium hydroxide. RNAs were again purified using MyOne Silane beads as described earlier. After drying, 2.5 µl of 5’ adapter master mix (4.66 µM InvRand3Tr3 adapter and 7.77% DMSO in TT elution buffer) were directly added to the beads and incubated at 70 °C for 2 min before 7.8 µl of ligation master mix (0.097% RNA ligase buffer (NEB), 1.94 mM DTT, 0.97 mM ATP, 0.019% v/v Tween-20, 17.48% PEG 8000, 30 U T4 RNA ligase high-concentration enzyme (NEB), 5’ deadenylase enzyme (NEB) and incubated at RT overnight. RNA was subsequently purified using MyONE silane beads as described earlier. To estimate the number of cycles required for the final PCR, qPCRs were conducted. cDNA libraries were amplified using a three-step, six cycle amplification followed by a two-step amplification bringing each sample to its calculated number of cycles. The final libraries were either purified using an agarose gel-based strategy cutting out gel pieces corresponding to 175-300 bp in size and purifying the DNA using the Qiagen MiniElute gel extraction kit following the manufacturer’s instructions. Alternatively, samples were cleaned using a two-sided selection with SPRI beads (0.7x right-sided, followed by a 1x left-sided selection). The quality and quantity of the libraries were assessed using the Fragment Analyzer (Agilent).

To account for different cycle number used during final PCR amplification (e.g. for IgG controls), a DNA spike-in containing adaptors for the Illumina index primers used in the final PCR was generated and equal amounts were added to the immunoprecipitation samples prior to the final PCR, resulting in different relative ratios of CLIP:DNA spike-in in the final libraries as observed upon measurement using the fragment analyzer.

### Multiplexed eCLIP sequencing

For multiplexing of eCLIPs, each RNA adapters containing unique barcodes (RNA_A01, RNA_B06, RNA_C01, RNA_D08, RNA_X1A, RNA_X1B) based on ^46^ were ligated to the RNA after immunoprecipitation. At least one third of each sample was mixed and further processed as a pool. SMI were separately run through the gel and barcoded RNA adapters were ligated after RNA isolation from the nitrocellulose membranes. As for the IP samples, inputs were merged to a pool and processed together. For total input samples, 20 µl of lysate were taken before the immunoprecipitation. Using the eCLIP adapter strategy ^46^, the in-line barcode on read 1 was later used to demultiplex the samples bioinformatically.

### DRIPc-Rx sequencing

DRIPc-sequencing was adapted from ^47^. Cells were lysed rotating overnight at 37 °C in TE buffer (pH 8) supplemented with 10% SDS and 10 µl proteinase K (10 mg/ml). For spike-normalization 10% *Drosophila* S2 cells were added according to the cell number. DNA was isolated by phenol/chloroform extraction using MaXtract High Density Tubes (Qiagen) and fragmented with a mixture of restriction enzymes. For this DNA was incubated overnight with 60 U of BsrGI, EcoRI, HindIII, SSpI and Xbal (New England Biolabs) at 37 °C. DNA was cleaned up again with phenol/chloroform and fragmentation was validated by agarose gel electrophoresis to ensure successful digestion of the DNA to a size below 1 kb. Subsequently, 200 µg of DNA was treated with 30 U of RNaseT1 (Thermo Fisher Scientific) at 37 °C for 30 min to eliminate ssRNA. To avoid potential dsRNA contamination the DNA was treated with 3 U RNaseIII (Thermo Fisher Scientific) for 1 h at 37 °C. Samples were then incubated for 6 h at 37 °C with or without 60 U of RNaseH1 (New England Biolabs) to ensure specificity of the following immunoprecipitation step. For this DNA in 1x DRIP binding buffer (10x DRIP binding buffer: 100 mM Sodium phosphate, 1.4 M NaCl, 0.5% Triton X-100) was split into two tubes and incubated overnight with 4 µl of S9.6 antibody (New England Biolabs) while rotating. For total input samples, 10% of DNA was taken before the immunoprecipitation. On the next day, 40 µl/IP of Protein A and Protein G-coupled Dynabeads (Thermo Fisher Scientific) were washed and added to the DNA/antibody mix and incubate for 4 h on a rotating wheel at 4 °C. Immunoprecipitated complexes were then washed 3 times each with ice-cold 1 x DRIP binding buffer. Finally, DNA was eluted by adding DRIP elution buffer (50 mM TRIS pH 8,10 mM EDTA, 0.5% SDS) and proteinase K for 45 min at 55 °C while shaking at 900 rpm. DNA was purified with phenol/chloroform extraction and finally eluted in 50 µl TE buffer. Before proceeding to library preparation, 4 µl were used to test enrichment by qPCR. DNA:RNA hybrids were then treated with DNase (Thermo Fisher Scientific) for 45 min at 37 °C. DNase was inactivated by inactivation buffer (Thermo Fisher Scientific) for 5 min at RT. After 5 min of centrifugation at 10,000 rpm the supernatant was transferred to a new PCR tube and libraries were generated using the NEBNext Ultra II Directional RNA Library Prep Kit for Illumina (New England Biolabs) following manufacturer’s instructions.

### mRNA sequencing

After treatment, cells were washed once with PBS and lysed with RLT buffer (QIAGEN) supplemented with beta-mercaptoethanol. Total RNA was extracted with RNeasy mini columns (QIAGEN) including an on-column DNase I digestion step according to manufacturer’s instructions. RNA concentration and RQN values were determined with a Fragment Analyzer (Agilent). RNA samples with RQN > 9 were used for mRNA isolation with the NEBNext Poly(A) mRNA Magnetic Isolation Module (New England Biolabs). cDNA libraries were then prepared using the NEBNext Ultra II Directional RNA Library Prep Kit for Illumina (New England Biolabs) according to manufacturer’s instructions. Libraries were amplified for 11 PCR cycles and size selected using SPRI selected beads (Beckman Coulter).

### Quantification and statistical analysis

#### ChIP-Rx data processing

Fastq files of ChIP-Rx samples were mapped separately to the human GRCh38 genome and to the murine GRCm39 genome using Bowtie1 or Bowtie2 with -N 1 parameters. Subsequently, aligned reads were spike-normalized by dividing the number of mapped reads aligned to GRCh38 by the number of reads mapped to GRCm39 for each sample and multiplying this ratio with the smallest number of reads aligned to GRCm39 for any sample.

### seCLIP data processing

For seCLIP experiments, the analysis was performed according to the Skipper workflow ^48^. For seCLIP experiments shown in Figures 2a,b,c,d, 3e, 4f and Extended Data Figure 2c,d,e, forward read (i5) fastq files of CLIP and input samples were normalized to sequencing depth using seqtk. Per condition, the same input subsampled using different random seeds was provided. For samples spiked with cDNA, reads mapping to the DNA spike-in were identified using STAR^49^. This applies to Figure 1b,c,d,e,f,g and Extended Data Figure 1a, b, c, d. Accordingly, fastq files were normalized according to the number of reads mapping to the spike-in in each sample. Multiplexed eCLIPs shown in Figure 2f,g and Extended Data Figures 2h, g, and 6c were demultiplexed using UMItools ^50^ and used without further normalization. Subsequently, samples were trimmed utilizing skewer ^51^ and extracts unimolecular identifiers (UMIs) using fastp ^52^ and aligned to the human genome version GRCh38 for all MYC seCLIP and eCLIP experiments or to the murine genome version GRCm38 for TLR3 seCLIPs using STAR ^49^. PCR duplicates were removed by using UMItools ^50^. To identify genomic regions with enriched signal in the CLIP over the SMI input, the automated Skipper analysis pipeline was used ^48^. The GENCODE version 38 gene annotations were obtained from gencodegenes.org. Transcripts were filtered based on total RNA-seq data in K562 cells with a minimum expression level of one transcript per million. A custom parse_gff.R script ^48^ was used to define RNA feature and transcript types (**Table 1**). Each feature was tiled into evenly spaced windows of maximum 100 nucleotides in length and then tested for enrichment in immunoprecipitated over SMI samples using a beta-binomial distribution that accounts for overdispersion in read counts and GC bias.

**Table 1:**
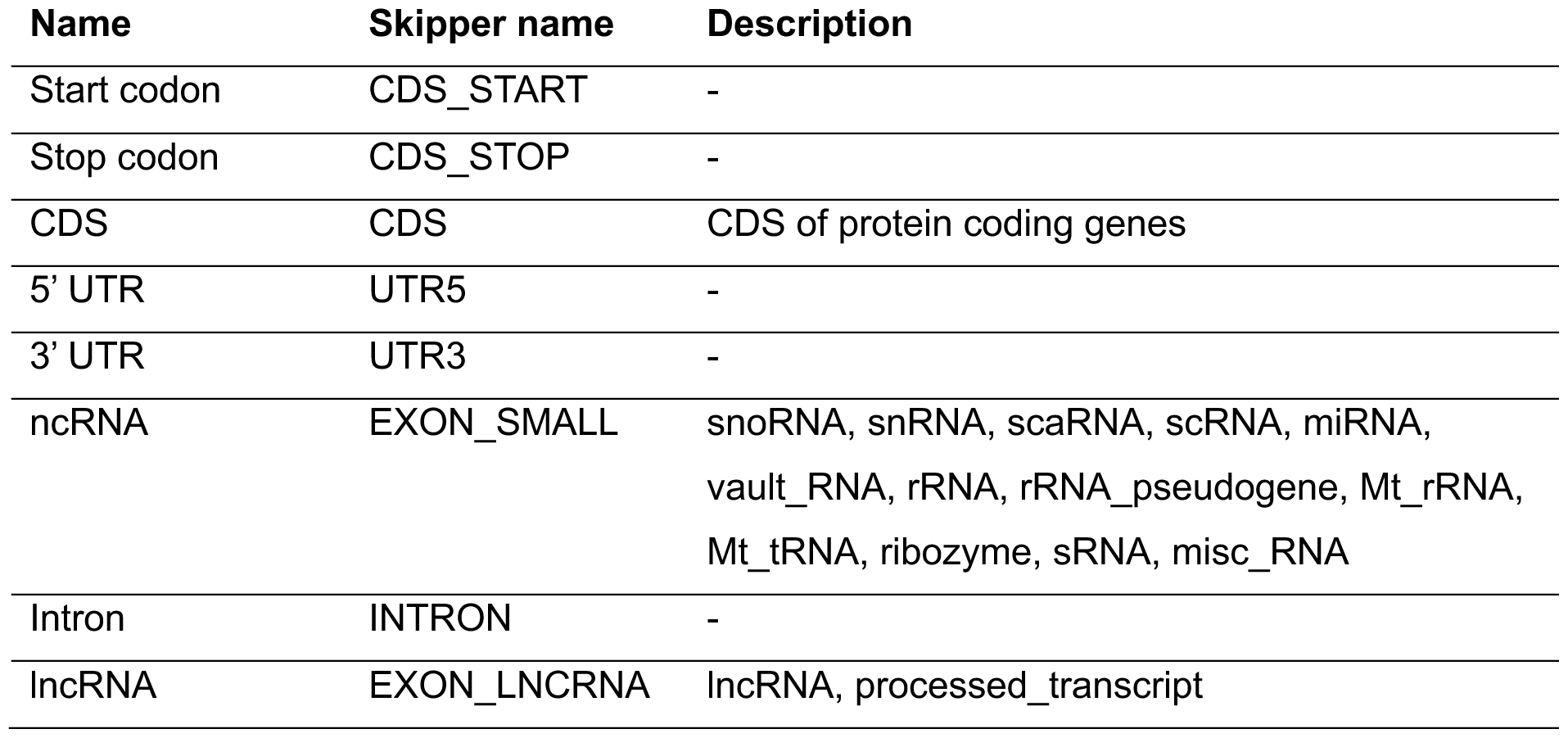
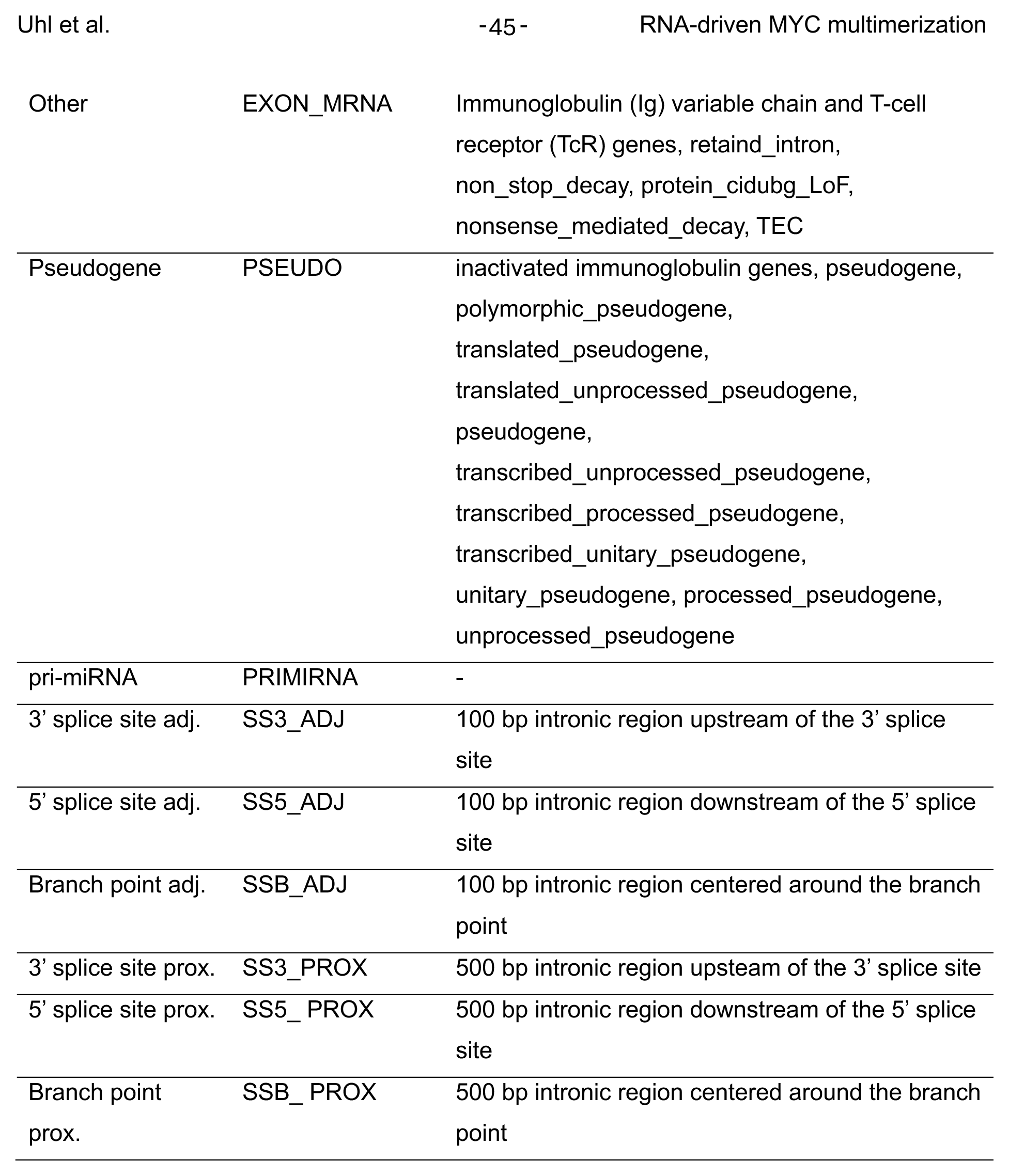
List of RNA feature types included in the Skipper output.

For the analysis of TLR3 seCLIPs by Skipper, statistical thresholds calculated from the fit of the beta-binomial distribution were derived from the automated Skipper pipeline (only windows passing a 20% false discovery rate in both replicates of each experiment were considered enriched), while for MYC seCLIP experiments the statistical thresholds were customized. Within individual replicates, windows that passed a *p* value threshold *p*<0.05 were considered enriched windows. Of these, windows that were identified in both replicates and passed a 20% FDR in at least one of the replicates were designated as reproducibly enriched windows. Windows that were called in more than 17% of all K562 and HepG2 ENCODE eCLIP samples were blacklisted and removed. For calling TLR3 reproducible enriched windows in the TLR3-IPs, reproducible enriched windows identified in IgG and NXL controls were blacklisted additionally. Reproducible windows were fine mapped by centering them around the local maximum of binding enrichment per window and adjusted to a fixed size of 75 nt. For RNA motif identification the HOMER script findMotifsGenome.pl (http://homer.ucsd.edu/homer/) was run with the following options: -preparsedDir <directory> -size given -rna -nofacts -S 20 - len 5,6,7,8,9 -nlen 1 -bg <background windows>. A random RNA background was specified by selecting random 75 nt windows belonging to the same RNA feature group. To visualize similarities in binding characteristics to different RNA transcript types in a t-SNE plot, performed MYC seCLIP data was pooled with ENCODE reference RBP eCLIP data, and plotted together in a two-dimensional space using the Rtsne package (https://lvdmaaten.github.io/tsne/). Several random seeds were tested to confirm a reproducible association of the queried protein with a certain feature type. Enrichment odds ratios (ORs), represented by the enrichment of observed CLIP enrichment over a baseline CLIP enrichment per RNA feature type, were calculated based on the call_enriched_windows.R script ^48^. For metagenes and heatmaps, log_2_FCs of CLIP/input were determined based on deduplicated bam files using bamCompare and plotted using deeptools ^53^ using the –sortUsingSamples and --averageTypeSummaryPlot mean options. Intersections between different sets of reproducible enriched windows were generated and visualized using intervene ^54^. Publicly available ENCODE eCLIP and control datasets were downloaded and log_2_ ratios of CLIP over SMIs were calculated using bamCompare and plotted over called windows of interest using deeptools^53^. For dot plots shown in Figure 6e and Extended Data Figure 6a, the RNA types were derived for the human (GRCh38) or the murine (GRCm38) genome using biomaRt ^55^. Mitochondrial (mt)RNA and ribosomal (r)RNA were excluded from the analysis.

### mRNA data processing

STAR (version: 2.7.10a) ^49^ was used to align the reads against the GRCh38 genome and quantify transcripts in the quant mode. The quantified transcripts were used as input for the R package DESeq2 ^56^ for differential expression analysis. Genes that passed FDR < 5% and had at least a 1.5-fold change in either direction with expression trend preserved across replicates were called significant. This resulted in 3,179 differentially expressed genes in shMYC vs ctrl and 2,428 in RBRIII^MUT^ vs WT MYC, with 2,126 overlapping genes between the two sets. DESeq2 was also used to normalize gene expression for downstream analysis. The heatmap was plotted with log_2_ normalized expression relative to average of the statistically significant genes using R package ComplexHeatmap ^57^. Genes from the MSigDB ^58^ pathways HALLMARK_MYC_TARGETS_V1, HALLMARK_MYC_TARGETS_V2, REACTOME_TRANSLATION, and HALLMARK_TNFA_SIGNALING_VIA_NFKB were annotated on the heatmap.

### DRIPc-Rx data processing

Fastq files of DRIPc-Rx samples were mapped separately to murine GRCm39 genome and to the fly dm6 genome using STAR ^49^. Subsequently, aligned reads were spike-normalized by dividing the number of mapped reads aligned to GRCm39 by the number of reads mapped to dm6 for each sample and multiplying this ratio with the smallest number of reads aligned to dm6 for any sample. Peaks were called using MACS ^59^with default broad peak settings. For metagenes and heatmaps, log_2_FC of DRIPs over inputs were determined based on bam files using bamCompare and plotted using deeptools ^53^.

### Statistical analysis of quantitative immunofluorescence

Quantitative immunofluorescence experiments were anaylzed using the Harmony High Content Imaging and Analysis Software (PerkinElmer) and single cell results were further processed in R v4.1.2. To identify MYC spots, the Find Spot building block was used and adjusted for each experiment based on corresponding positive and negative controls. To calculate *PCCs*, the MYC partitioning ratio (average MYC intensity in the MYC spot/average MYC intensity in the nucleus) and the equally calculated partitioning ratio of the co-stained protein across all spots was determined. For each condition more than 500 cells were used. For plots shown in figure 3g and 5f, single cells were binned based on their MYC nuclear intensities and the average number of spots per bin were calculated. Dose-response modelling was used to evaluate the relationship between average MYC intensity per bin and average number of spots per bin in different samples and the R packages drc and ggplot2 were used for model fit and visualization respectively. Two-sample *t*-test was used to determine statistical significance between samples with 4 or 8 replicates each.

For analysis of siRNA-based screens presented in figure 4a and c, unpaired two-sided *t*-tests were performed on at least three independent experiments, and data are presented as mean ± s.e.m. Data points of individual replicates are indicated as data points.

## Data availability

RNA-, DRIPc-, ChIP-Rx-, seCLIP-, datasets are being submitted and will be available at the Gene Expression Omnibus under the accession GSE281781.

## Acknowledgements

This work was supported by grants from the European Research Council (SENATR ERC #101096948 to M.E.), the German Cancer Aid (#70114538 to M.E.), the Mildred Scheel Junior Research Center Program (to D.P.), the German Research Foundation (MA6957/1-1 to H.M.M., EI 222/21-1 and INST 93/1023-1-FUGG to M.E.), the Cancer Research UK (to M.E., S.M.V. and T.R.) and the Alex’s Lemonade Foundation Crazy 8 Initiative (to S.M.V. and M.E.). S.M.V. is a Freeman Hrabowksi Scholar of the Howard Hughes Medical Institute. The authors thank Carsten Ade for performing the sequencing experiments, Peter Gallant for help with bioinformatic analyses and Tobias Roth and Ciler Maden for technical support.

## Author information

These authors contributed equally: Leonie Uhl, Amel Aziba.

## Contributions

L.U., A.A., S.L., T.M., M.B. and T.E. performed most of the experiments. L.U., T.K., F.M., D.S. and D.P. performed bioinformatic anaylsis. O.V. performed peptide library synthesis, microarray printing and binding assay. T.R. performed purification of MYC constructs fluorescence anisotropy assays. B.K. and G.C. provided unpublished data. C.S.-V. performed high-content immunofluorescence experiments. H.M.M., D.P., S.M.V. and M.E. devised and supervised experiments and M.E. wrote the paper.

## References

1. Dang, C.V. MYC on the path to cancer. Cell 149, 22–35 (2012).

2. Kress, T.R., Sabo, A. & Amati, B. MYC: connecting selective transcriptional control to global RNA production. Nat Rev Cancer 15, 593–607 (2015).

3. Dhanasekaran, R. et al. The MYC oncogene - the grand orchestrator of cancer growth and immune evasion. Nature reviews. Clinical oncology 19, 23–36 (2022).

4. Muthalagu, N. et al. Repression of the Type I Interferon pathway underlies MYC & KRAS-dependent evasion of NK & B cells in Pancreatic Ductal Adenocarcinoma. Cancer Discov (2020).

5. Zimmerli, D. et al. MYC promotes immune-suppression in triple-negative breast cancer via inhibition of interferon signaling. Nat Commun 13, 6579 (2022).

6. Kortlever, R.M. et al. Myc Cooperates with Ras by Programming Inflammation and Immune Suppression. Cell 171, 1301–1315 e1314 (2017).

7. Dhanasekaran, R. et al. MYC Overexpression Drives Immune Evasion in Hepatocellular Carcinoma That Is Reversible through Restoration of Proinflammatory Macrophages. Cancer research 83, 626–640 (2023).

8. Krenz, B. et al. MYC- and MIZ1-Dependent Vesicular Transport of Double-Strand RNA Controls Immune Evasion in Pancreatic Ductal Adenocarcinoma. Cancer research 81, 4242–4256 (2021).

9. Gaballa, A. et al. PAF1c links S-phase progression to immune evasion and MYC function in pancreatic carcinoma. Nat Commun 15, 1446 (2024).

10. Bernards, R., Dessain, S.K. & Weinberg, R.A. N-myc amplification causes down-modulation of MHC class I antigen expression in neuroblastoma. Cell 47, 667–674 (1986).

11. Runde, A.P., Mack, R., S, J.P. & Zhang, J. The role of TBK1 in cancer pathogenesis and anticancer immunity. Journal of experimental & clinical cancer research : CR 41, 135 (2022).

12. Zheng, X., Li, S. & Yang, H. Roles of Toll-Like Receptor 3 in Human Tumors. Frontiers in immunology 12, 667454 (2021).

13. Crossley, M.P. et al. R-loop-derived cytoplasmic RNA-DNA hybrids activate an immune response. Nature 613, 187–194 (2023).

14. Solvie, D. et al. MYC multimers shield stalled replication forks from RNA polymerase. Nature 612, 148–155 (2022).

15. Yang, J. et al. MYC phase separation selectively modulates the transcriptome. Nat Struct Mol Biol 31, 1567–1579 (2024).

16. Papadopoulos, D. et al. The MYCN oncoprotein is an RNA-binding accessory factor of the nuclear exosome targeting complex. Mol Cell 84, 2070–2086 e2020 (2024).

17. Oksuz, O. et al. Transcription factors interact with RNA to regulate genes. Mol Cell 83, 2449–2463 e2413 (2023).

18. Muhar, M. et al. SLAM-seq defines direct gene-regulatory functions of the BRD4-MYC axis. Science 360, 800–805 (2018).

19. Blue, S.M. et al. Transcriptome-wide identification of RNA-binding protein binding sites using seCLIP-seq. Nature protocols 17, 1223–1265 (2022).

20. Xu, S. et al. Protocol to process crosslinking and immunoprecipitation data into annotated binding sites. STAR Protoc 5, 103040 (2024).

21. Uemura, Y. et al. Matrin3 binds directly to intronic pyrimidine-rich sequences and controls alternative splicing. Genes Cells 22, 785–798 (2017).

22. Attig, J. et al. Heteromeric RNP Assembly at LINEs Controls Lineage-Specific RNA Processing. Cell 174, 1067–1081 e1017 (2018).

23. Coelho, M.B. et al. Nuclear matrix protein Matrin3 regulates alternative splicing and forms overlapping regulatory networks with PTB. The EMBO journal 34, 653–668 (2015).

24. Ray, D. et al. A compendium of RNA-binding motifs for decoding gene regulation. Nature 499, 172–177 (2013).

25. Lorenzin, F. et al. Different promoter affinities account for specificity in MYC-dependent gene regulation. eLife 5 (2016).

26. Lin, C.Y. et al. Transcriptional amplification in tumor cells with elevated c-Myc. Cell 151, 56–67 (2012).

27. Kaida, D. et al. Spliceostatin A targets SF3b and inhibits both splicing and nuclear retention of pre-mRNA. Nat Chem Biol 3, 576–583 (2007).

28. Kotake, Y. et al. Splicing factor SF3b as a target of the antitumor natural product pladienolide. Nat Chem Biol 3, 570–575 (2007).

29. Gockert, M. et al. Rapid factor depletion highlights intricacies of nucleoplasmic RNA degradation. Nucleic Acids Res 50, 1583–1600 (2022).

30. Lykke-Andersen, S., Rouviere, J.O. & Jensen, T.H. ARS2/SRRT: at the nexus of RNA polymerase II transcription, transcript maturation and quality control. Biochem Soc Trans 49, 1325–1336 (2021).

31. Thomas, L.R. et al. Interaction of MYC with host cell factor-1 is mediated by the evolutionarily conserved Myc box IV motif. Oncogene 35, 3613–3618 (2016).

32. Nair, S.K. & Burley, S.K. X-ray structures of Myc-Max and Mad-Max recognizing DNA. Molecular bases of regulation by proto-oncogenic transcription factors. Cell 112, 193–205 (2003).

33. Dang, C.V. & Lee, W.M.F. Identification of the human c-myc protein nuclear translocation signal. Mol. Cell. Biol. 8, 4048–4054 (1988).

34. Hingorani, S.R. et al. Trp53R172H and KrasG12D cooperate to promote chromosomal instability and widely metastatic pancreatic ductal adenocarcinoma in mice. Cancer cell 7, 469–483 (2005).

35. Alexopoulou, L., Holt, A.C., Medzhitov, R. & Flavell, R.A. Recognition of double-stranded RNA and activation of NF-kappaB by Toll-like receptor 3. Nature 413, 732–738 (2001).

36. Crossley, M.P. et al. R-loop-derived cytoplasmic RNA-DNA hybrids activate an immune response. Nature 613, 187–194 (2023).

37. Webster, S.F. & Ghalei, H. Maturation of small nucleolar RNAs: from production to function. RNA biology 20, 715–736 (2023).

38. Larochelle, S. et al. Cyclin-dependent kinase control of the initiation-to-elongation switch of RNA polymerase II. Nat Struct Mol Biol 19, 1108–1115 (2012).

39. Olson, C.M. et al. Pharmacological perturbation of CDK9 using selective CDK9 inhibition or degradation. Nat Chem Biol 14, 163–170 (2018).

40. Yang, J. et al. MYC phase separation selectively modulates the transcriptome. Nat Struct Mol Biol (2024).

41. Boija, A. et al. Transcription Factors Activate Genes through the Phase-Separation Capacity of Their Activation Domains. Cell 175, 1842–1855 e1816 (2018).

42. Wadsworth, G.M. et al. RNA-driven phase transitions in biomolecular condensates. Mol Cell 84, 3692–3705 (2024).

43. Yu, Y.T. & Meier, U.T. RNA-guided isomerization of uridine to pseudouridine--pseudouridylation. RNA biology 11, 1483–1494 (2014).

44. Berndt, H. et al. Maturation of mammalian H/ACA box snoRNAs: PAPD5-dependent adenylation and PARN-dependent trimming. Rna 18, 958–972 (2012).

45. Schulte, C., Khayenko, V. & Maric, H.M. Peptide Microarray-Based Protein Interaction Studies Across Affinity Ranges: Enzyme Stalling, Cross-Linking, Depletion, and Neutralization. Methods Mol Biol 2578, 143–159 (2023).

46. Van Nostrand, E.L. et al. Robust transcriptome-wide discovery of RNA-binding protein binding sites with enhanced CLIP (eCLIP). Nature methods 13, 508–514 (2016).

47. Sanz, L.A. & Chedin, F. High-resolution, strand-specific R-loop mapping via S9.6-based DNA-RNA immunoprecipitation and high-throughput sequencing. Nature protocols 14, 1734–1755 (2019).

48. Boyle, E.A. et al. Skipper analysis of eCLIP datasets enables sensitive detection of constrained translation factor binding sites. Cell Genom 3, 100317 (2023).

49. Dobin, A. et al. STAR: ultrafast universal RNA-seq aligner. Bioinformatics 29, 15–21 (2013).

50. Smith, T., Heger, A. & Sudbery, I. UMI-tools: modeling sequencing errors in Unique Molecular Identifiers to improve quantification accuracy. Genome Res 27, 491–499 (2017).

51. Jiang, H., Lei, R., Ding, S.W. & Zhu, S. Skewer: a fast and accurate adapter trimmer for next-generation sequencing paired-end reads. BMC Bioinformatics 15, 182 (2014).

52. Chen, S., Zhou, Y., Chen, Y. & Gu, J. fastp: an ultra-fast all-in-one FASTQ preprocessor. Bioinformatics 34, i884–i890 (2018).

53. Ramirez, F., Dundar, F., Diehl, S., Gruning, B.A. & Manke, T. deepTools: a flexible platform for exploring deep-sequencing data. Nucleic Acids Res 42, W187–191 (2014).

54. Khan, A. & Mathelier, A. Intervene: a tool for intersection and visualization of multiple gene or genomic region sets. BMC Bioinformatics 18, 287 (2017).

55. Durinck, S., Spellman, P.T., Birney, E. & Huber, W. Mapping identifiers for the integration of genomic datasets with the R/Bioconductor package biomaRt. Nature protocols 4, 1184–1191 (2009).

56. Love, M.I., Huber, W. & Anders, S. Moderated estimation of fold change and dispersion for RNA-seq data with DESeq2. Genome Biol 15, 550 (2014).

57. Gu, Z. Complex heatmap visualization. Imeta 1, e43 (2022).

58. Liberzon, A. et al. Molecular signatures database (MSigDB) 3.0. Bioinformatics 27, 1739–1740 (2011).

59. Zhang, Y. et al. Model-based analysis of ChIP-Seq (MACS). Genome Biol 9, R137 (2008).

